# Structural Maintenance of Chromosome 3 interacts with the Topoisomerase VI complex and contributes to the oxidative stress response in *Arabidopsis thaliana*

**DOI:** 10.1101/2022.10.27.514040

**Authors:** Florent Velay, Dina Abdallah, Cécile Lecampion, Nadia Kbiri, Stefano D’Alessandro, Benjamin Field, Christophe Laloi

## Abstract

In plants adverse environmental conditions can induce the accumulation of reactive oxygen species, such as singlet oxygen or hydrogen peroxide, at the level of the photosynthetic apparatus. The coordinated action of nucleus-encoded genes is required for containing the deleterious effects of reactive oxygen species. The regulation of such genes follows a molecular signalling process between the chloroplast and the nucleus called retrograde signalling. Previously, we proposed that the Topoisomerase VI (Topo VI) complex participates in the singlet oxygen stress response by regulating the expression of specific subsets of nuclear genes. However, the underlying molecular mechanisms remain unresolved. In this study, we demonstrate that the Topo VI subunit BIN4 interacts with the cohesin subunit AtSMC3. We also show that, similarly to Topo VI mutants, a line suppressing AtSMC3 shows constitutive activation of singlet oxygen response genes and enhanced tolerance to photooxidative stress. Together, these results suggest that Topo VI and AtSMC3 control the expression of singlet oxygen response genes and are possibly involved in the acclimation of plants to photooxidative stress conditions.

## INTRODUCTION

Abrupt changes in environmental conditions are a source of stress that can disrupt cellular homeostasis and affect the integrity of organisms. One of the consequences common to most environmental stresses is the production of reactive oxygen species (ROS), such as hydrogen peroxide (H_2_O_2_) or singlet oxygen (^1^O_2_) (Baxter, Mittler and Suzuki, 2014). Although the oxidizing power of ROS can be deleterious for the function of a wide range of macromolecules, they can also act as signals and promote the induction of several stress responsive genes, at lower concentrations (Laloi and Havaux, 2015; Exposito-Rodriguez *et al*., 2017). In plants, ROS accumulation arise mainly from an imbalance between energy harvesting and dissipation at the level of the photosynthetic chain in the chloroplasts (Pinnola and Bassi, 2018). Therefore, ROS signalling is among the principal actor of inter-organellar communication from chloroplasts to the nucleus, namely retrograde signal.

The less reactive H_2_O_2_ has been shown to diffuse out of isolated chloroplasts, probably through aquaporins (Mubarakshina *et al*., 2010; Bienert and Chaumont, 2014). Furthermore, direct transfers of H_2_O_2_ from chloroplasts to nucleus have been highlighted in *Nicotiana benthamiana* during high light stress, followed by an activation of *NbAPXa*, a H_2_O_2_ responsive gene, suggesting that H_2_O_2_ could act as a signal itself (Exposito-Rodriguez *et al*., 2017). Because of its short lifetime and high reactivity, ^1^O_2_ is unlikely to follow the same signalling scheme as H_2_O_2_. In the presence of ^1^O_2_, induction of cell death can be mediated by two independent pathways involving EXECUTERs or OXI1 proteins (Laloi and Havaux, 2015). Moreover, the activation of photooxidative stress tolerance nuclear genes rather relies on the production of β-cyclocitral (β-CC), a product of the oxidation of β-carotene (Ramel *et al*., 2012). Following this, METHYLENE-BLUE SENSITIVITY 1 (MBS1) and SCARCROW LIKE14 (SCL14), two transcriptional regulators induced by β-CC, regulate the acclimation process through two independent pathways (Shumbe *et al*., 2017; D’alessandro, Ksas and Havaux, 2018; Dmitrieva, Tyutereva and Voitsekhovskaja, 2020). The action of such transcriptional regulators on the nuclear genome is intrinsically regulated by chromatin topology. Indeed, studies describing the involvement of chromatin remodelling complexes and topoisomerases in stress response are increasing (Vriet, Hennig and Laloi, 2015; Song, Liu and Han, 2021), thus revealing the importance of chromatin architecture regulation in the completion of retrograde signalling.

To identify ^1^O_2_ dependant retrograde signalling compounds, we previously performed a genetic screen that allowed the isolation of constitutive activators (*caa*) as well as non-activators (*naa*) of the *AAA-ATPase (AAA)* / *AT3G28580*, a ^1^O_2_-responsive gene (Baruah *et al*., 2009). We reported that a weak mutant allele of the A subunit of the topoisomerase VI (Topo VI) complex, *caa39*, constitutively activates a transcriptional response to ^1^O_2_ that cannot be further enhanced under stress conditions (Šimková *et al*., 2012). Moreover, chromatin immunoprecipitation experiments showed that two subunits of the Topoisomerase VI (TopoVIA and RHL1) bind the proximal regions of ^1^O_2_-responsive gene promoters during high light stress (Šimková *et al*., 2012). Interestingly, H_2_O_2_ response genes are stimulated in the *caa39* mutant under photooxidative stress (Šimková *et al*., 2012). Considering the antagonistic effects of the *caa39* mutation on H_2_O_2_ and ^1^O_2_ pathways under photoinhibitory conditions, we proposed Topo VI to be a molecular switch that might relay both H_2_O_2_ and ^1^O_2_ responses. To date, the mechanistic insights of the Topo VI-dependent retrograde signalling and its connexion with the above-mentioned retrograde signalling components have not been completely elucidated.

In this study, we show that the Topo VI subunit BIN4 interacts with the cohesin subunit SMC3, likely through the hinge domain of SMC3. Since knockout of *SMC3* is embryonically lethal (Schubert *et al*., 2009), we isolated *SMC3* co-suppression lines to understand the genetic interaction between Topo VI and SMC3. The transcriptomic analysis of *SMC3* co-suppression lines and *caa39* mutants revealed that both lines constitutively activate ^1^O_2_-responsive genes. Finally, we show that *caa39* and *SMC3* co-suppression lines display enhanced resistance to high light stress conditions, which is correlated with the activation of the non-photochemical quenching mechanisms.

## MATERIALS AND METHODS

### Plant material and growth conditions

*Nicotiana benthamiana* was used for transient expression of recombinant proteins. *caa39* mutant and SMC3 suppressors are in *Arabidopsis thaliana* Col-0 ecotype. Both Arabidopsis and Nicotiana plants were grown in a controlled environment at 120 μmol photons m^-2^ s^-1^ illumination with an 8 h (short days) or 16 h (long days) photoperiod at 22°C day / 20°C night, and 55% day / 75% night relative humidity. The photoperiod used for each experiment is indicated in the figure legends.

### Cloning

Multigenic BiFC vectors containing a repressor of silencing (P19), a transformation marker (OEP7-mTRQ), and the two coding sequences of interest fused with the two different parts of the 174/175 YFP split were made using the cloning toolbox and the protocol described in (Engler *et al*., 2014; Velay *et al*., 2022). The chimeric CDS of *AtSMC3* was cloned following an optimized protocol due to the high instability of the construct in bacterial strains. Notably, the addition of genomic introns n°1, 24, 25, 26, 27 was found to help improve stability. Reducing the number of bacteria spread on selective media (maximum 10 colonies per Petri dish) also helped to dramatically reduce the frequency of recombination events within the *AtSMC3* CDS. Moreover, due to the strong propensity of positive colonies grown on solid media to quickly evict the *AtSMC3*-containing plasmid, after each cloning step, the positive clones were constantly kept in a liquid culture containing the appropriate antibiotic and refreshed every 4 days. After transformation of the final plasmid into *Agrobacterium tumefaciens*, no notable instability was detected.

### Yeast two-hybrid screen

The yeast two-hybrid screen was performed by Hybrigenics using the Arabidopsis RP1 library. The full– length *BIN4* cDNA (*AT5G24630*.*3/4*) was used as bait.

### Stable and transient expression by agroinfiltration

Transient expression in *Nicotiana benthamiana* leaves was performed following the protocol described in Velay *et al*. 2022. For Arabidopsis stable expression, *Agrobacterium tumefaciens* GV3101 transformed with the plasmid containing GFP-SMC3 were grown at 28°C in LB medium supplemented with rifampicin and kanamycin. Arabidopsis flowers were then dipped in 50 mL of Agrobacterium suspension OD_600_ = 0.8 containing 5 % sucrose and 0.05 % SILWET L-77 (Clough and Bent, 1998). The transformed plants were covered and kept in the dark for 24 h after what they were manually watered every 4 days until the seeds were harvested. Bialaphos resistant primary transformants were selected by spraying a solution containing 150 mg/L ammonium-glufosinate on 6-day-old seedlings.

### BiFC assay

BiFC assays were carried out in *Nicotiana benthamiana* lower epidermal cells. Sample preparation, imaging and analysis were performed following the protocol described in (Velay *et al*., 2022). Exposure time and post-acquisition analyses were similar between all samples.

### Protein extraction, antibody production and Immunoblotting

For total protein extraction, 6-day-old plant aerial parts; 2-week-old plant aerial parts; 4-week-old plant mature rosette leaves were harvested, and ground in liquid nitrogen. Tissue powder was resuspended in SDS loading buffer, heated at 85°C 10 min and centrifuged at 16000 g, 10 min at room temperature. The proteins contained in the supernatant were then separated by SDS-PAGE, transferred onto a nitrocellulose membrane and probed with specific antibodies. To generate the anti-AtSMC3 antibody, we synthesized the peptide C-QALDFIEKDQSHDT, corresponding to the C-terminal region of AtSMC3, and used it to raise a polyclonal antibody in rabbit (Genscript). Anti-AtSMC3 and anti-GFP (Roche 11814460001) antibodies were diluted 1:500 and 1:1000 in PBST milk 5 %, respectively. Total protein staining was performed using SYPRO ruby protein stain (Thermofisher).

### RNA extraction and RT-qPCR

Total RNA was extracted using TRI Reagent (MRC). 500 ng of RNA was treated with DNaseI (Takara) and then used for RT reactions using the PrimeScript RT Reagent Kit (Takara), with an equimolar mix of random hexamers and polyA primers. The efficiency of RT reaction and absence of residual genomic DNA were confirmed by semi-quantitative RT-PCR. cDNAs were then diluted 4 times with ultra-pure water. Quantitative RT-PCR were carried out using TB Green Premix Ex Taq II (Takara). 1 μL of diluted cDNA was used for each reaction in a final volume of 15 μL. *PP2A* (*AT1G13320*) and *PRF1* (*AT2G19760*) were used as control genes in all RT-qPCR experiments.

### RNA-seq library preparation and sequencing

Two independent biological replicates were produced. For each biological repetition, RNA samples were obtained by pooling RNAs from 4 plants. Aerials parts were collected on plants at developmental growth stage 1.04 (Boyes *et al*., 2001) cultivated as described above. Total RNA was extracted using TRI Reagent (MRC) and purified using RNA clean and concentrator-25 (Zymo Research) according to the supplier’s instructions. Libraries were prepared and sequenced at BGI (China) on a DNBSEQ™ sequencing platform in paired-end (PE) with a read length of 100 bases. Approximately 30 million of PE reads by sample were generated.

### RNA-seq bioinformatic treatment and analyses

Each RNA-Seq sample was analysed using the same workflow. Read preprocessing criteria were assessed using FastQC (v0.11.9). Bowtie 2 (v 2.4.4) (Langmead and Salzberg, 2012) was used to align reads against the *Arabidopsis thaliana* transcriptome (Araport11_cdna_20160703_representative_gene_model.fa). The reads count was calculated by a local script adapted from Van Verk *et al*., 2013 (Van Verk *et al*., 2013). Gene expression analysis was performed using SarTools (v1.7.4) (Varet *et al*., 2016) with EdgeR in R (v4.1.2) (Robinson, McCarthy and Smyth, 2009). Cluster analysis was performed using a script derived from https://2-bitbio.com/2017/10/clustering-rnaseq-data-using-k-means.html (Method S1). The optimal number of clusters was calculated using four different methods: sum os squared error, average silhouette width, Calinsky criterion and gap statistic (Calinski and Harabasz, 1974; Rousseeuw, 1987; Tibshirani, Walther and Hastie, 2001). GO enrichment analysis of the RNA-seq data was performed using a custom script (prepare_gene_ontology.pl, https://github.com/cecile-lecampion/gene-ontology-analysis-and-graph) which automatically uses PANTHER and REVIGO for the identification and simplification of enriched GO terms according to the procedure proposed by Bonnot *et al*., 2019 (Bonnot, Gillard and Nagel, 2019). Results were plotted using ggplot2.

### High light stress and photosynthetic parameter measurements

2-week or 4-week-old plants grown in short-day conditions were submitted to two high light (HL) stress periods consisting of 1500 μmol photons m^-2^ s^-1^ illumination at 20°C with a 8 h light / 16 h dark photoperiod (Ramel *et al*., 2012). Plants were mounted on a grill to allow air circulation at the base of plants. Photosynthetic parameters were acquired on dark adapted plants using a FluorCam (Photon System Instruments) before HL, after HL and after 16 h of recovery period in the dark. Plants were rewatered at the beginning of the recovery period.

### Data analysis

Graphs and statistical tests were generated in Python (Python Software Foundation, https://www.python.org/) using the Panda (Reback *et al*., 2022), Matplotlib (Hunter, 2007), Seaborn (Waskom, 2021) and Pingouin (Vallat, 2018) libraries. A Games-Howell Post-hoc test was adopted for non-parametric data comparisons and a pairwise T-test for data with normal distributions. For multiple comparisons, Games-Howell Post-hoc test was adopted for non-parametric data and a pairwise T-test using the Benjamini/Hochberg FDR correction for data with normal distributions.

## RESULTS

### The Topoisomerase VI subunit BIN4 interacts with SMC3

To gain insight into the molecular mechanism by which Topo VI controls the expression of oxidative stress responsive genes, we performed a yeast two hybrid (Y2H) screen with the BIN4 subunit as bait. The screen was performed under two different stringency conditions, 2.0 and 0.5 mM 3AT (Hybrigenics, Table S1). We detected a strong interaction with the Topo VI subunit RHL1, which supports the reliability of the screening procedure (Table S1). Among the eleven additional interacting partners, TTN7 (TITAN7) / SMC3 (STRUCTURAL MAINTENANCE OF CHROMOSOMES 3) / AT2G27170 was identified under the two stringency conditions (Table S1). *AtSMC3* encodes a subunit of cohesin, a ring shape chromatin architectural protein complex involved in sister chromatid cohesion (Tao *et al*., 2007; Schubert *et al*., 2009; Bolaños-Villegas *et al*., 2013; Morales and Losada, 2018), gene expression (Dorsett and Merkenschlager, 2013), DNA repair (Sjögren and Ström, 2010; Bolaños-Villegas *et al*., 2013; da Costa-Nunes *et al*., 2014; Pradillo *et al*., 2015), loop extrusion (Rowley and Corces, 2018) and many other processes related to chromatin dynamics (Cheng, Zhang and Pati, 2020; Bolaños-Villegas, 2021). The AtSMC3 protein is composed of two globular domains located at the C- and N-terminal extremities, forming an ATPase head necessary for the proper conformation of the cohesin ring during DNA entrapment or release (Arumugam *et al*., 2003; Murayama and Uhlmann, 2014; Muir *et al*., 2020). The overall rod shape structure of AtSMC3 is due to the presence of two coil domains, separated by a hinge, having the capacity to associate to each other, forming a coiled-coil region (Haering *et al*., 2002; Jumper *et al*., 2021). Among the two clones isolated in Y2H, the clone pB27A-36 encodes a truncated N-terminal coil, the entire hinge domain, and a truncated C-terminal coil (SMC3 193-807) meanwhile the second clone would only encode a truncated hinge domain and a truncated C-terminal coil.

We then sought to confirm the biochemical interaction between AtSMC3 and BIN4 *in planta* using Bimolecular Fluorescence Complementation (BiFC). We designed two recombinant CDSs, BIN4 fused with the C-terminus of YFP as well as the CDS of SMC3(193-807) fused with the N-terminus of YFP and supplemented with a Simian-Virus 40 Nuclear Localization Signal (NLS) to ensure the nuclear location of the truncated SMC3 peptide. We then added both CDS of interest to the same multigenic expression vector that also contains the silencing repressor P19 and the chloroplast outer membrane OEP7-mTRQ as a transformation marker (Yong Jik Lee *et al*., 2001; Scholthof, 2006; Velay *et al*., 2022). Expression of BIN4-cYFP and SMC3(193-807)-nYFP resulted in reconstitution of YFP fluorescence in the nuclei of epidermal cells of *Nicotiana benthamiana* (Fig. 1). To determine which part of SMC3 is necessary for the interaction, we designed 3 truncated versions of SMC3(193-807): SMC3(602-807), lacking the N-terminal coil and the N-terminal half of the hinge domain; SMC3(193-555), lacking the C-terminal coil and the C-terminal half of the hinge domain; SMC3(193-807, Δ525-632) lacking the entire hinge domain (Fig. 1a). Expression of SMC3(602-807)-nYFP with BIN4-cYFP led to a significant drop in YFP fluorescence compared to experiments using SMC3(193-807)-nYFP (Fig. 1b,c). The use of SMC3(193-555)-nYFP resulted in an even greater decrease in YFP fluorescence. Finally, the SMC3(193-807, Δ525-632)-nYFP/BIN4-cYFP pair produced only background fluorescence in the transformed nuclei (Fig. 1b,c). Collectively, these results show that BIN4 and SMC3 can interact *in planta* and that the hinge domain of SMC3 is necessary for this interaction.

**Fig. 1.**
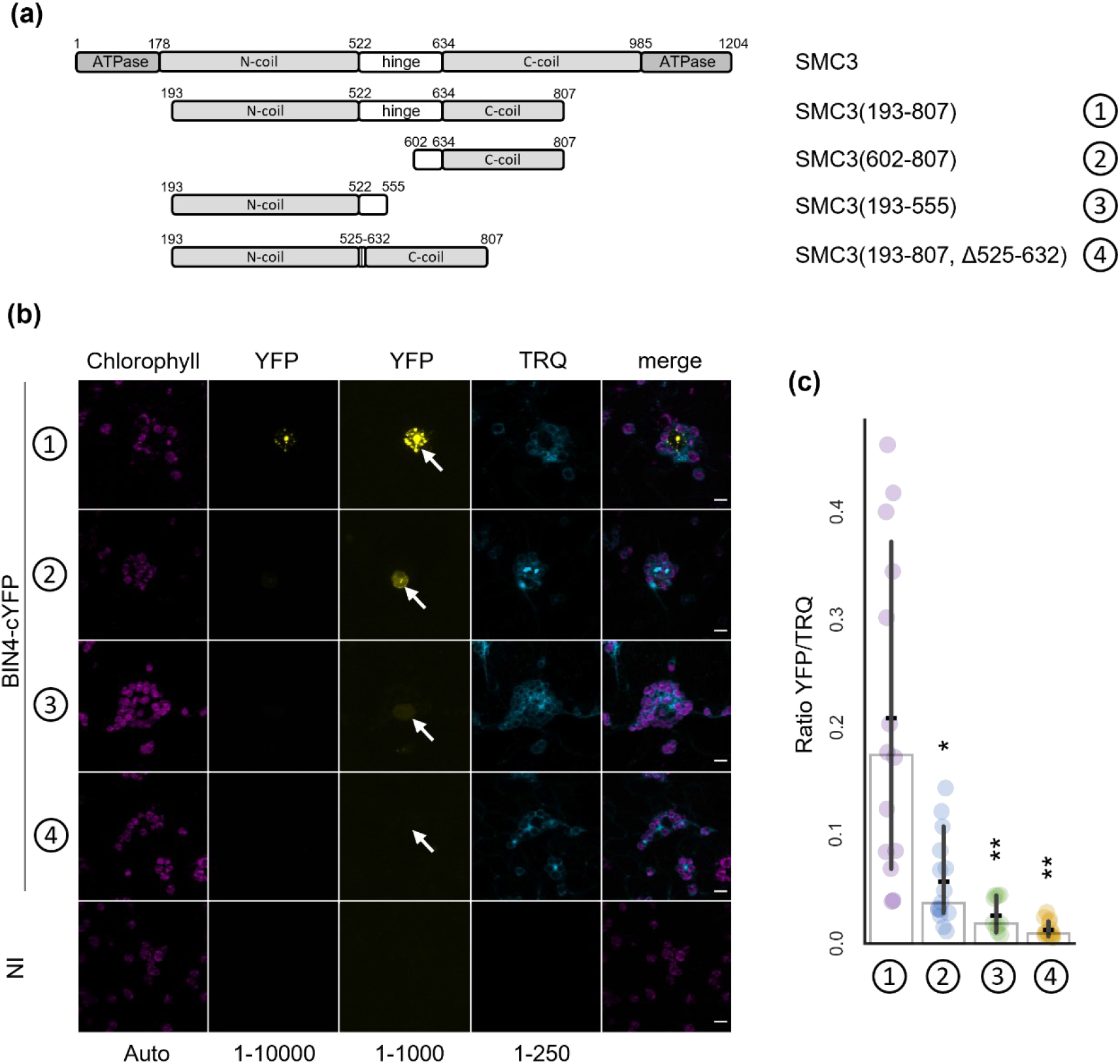
SMC3 interacts with BIN4 *in planta*. (a) Schematic representation of the different domains of AtSMC3; (1) SMC3(193-807); (2) SMC3(602-807); (3) SMC3(193-555); (4) SMC3(193-807, Δ525-632). (b) BiFC assay in *Nicotiana benthamiana* epidermal cells showing (1) the nuclear interaction of BIN4-cYFP and NLS-SMC3(193-807)-nYFP; (2) the weak interaction of BIN4-cYFP and NLS-SMC3(602-807)-nYFP; (3) the weaker interaction of BIN4-cYFP and NLS-SMC3(193-555)-nYFP; (4) the absence of interaction between BIN4-cYFP and NLS-SMC3(193-807, Δ525-632)-nYFP. The two YFP columns correspond to two distinct histogram levels. Histogram levels are indicated below each column. The positions of the nuclei are indicated by the white arrows. NI, not inoculated; scale bar, 10 μm. (c) Normalized BiFC signal consisting of the ratio between the YFP and the TRQ fluorescences. (1), (2), (3) and (4) as in (b) and (c). Bars indicate mean and crosses indicate median +/-95% confidence interval (n=10-15 nuclei). Statistical tests (Games-Howell Post-hoc test) shown against (1), * *P<0,05*, ** *P<0,01*. The whole experiment was repeated twice with similar results.

### Generation and characterization of *35S::GFP-SMC3* transgenic lines

We initially aimed to generate transgenic Arabidopsis plants overexpressing tagged SMC3 to allow biochemical studies of SMC3. For this purpose, we cloned the full CDS of *AtSMC3* following an optimized protocol due to the high instability of the construct in bacterial strains. Notably, the addition of genomic introns n°1, 24, 25, 26, 27 was found to help improve stability. We transformed *Arabidopsis thaliana* plants with the synthetic transcription unit *35S::GFP-SMC3* associated with a Bialaphos Resistance (BaR) cassette. Seven out of sixteen independent T1 plants showed a similar morphological phenotype with varying degrees of severity among individuals: reduced growth, short petioles, anthocyanin accumulation and reduced fertility. The most severely affected plants were sterile (Fig. S1a). Based on the BaR segregation in the T2 generations from the fertile T1 plants, we selected four independent transformant lines likely containing a single T-DNA insertion. Similarly, we then isolated homozygous individuals based on BaR segregation in the T3s. Among the four lines isolated this way, three displayed an almost identical morphological phenotype characterized by reduced growth, short petioles, anthocyanin accumulation, reduced fertility and, after 5 weeks, leaf necrotic spots, reminiscent of hypersensitive response (Fig. S1b). The fourth line displayed a less severe morphological phenotype only visible after 4 weeks of growth. Suspecting that the phenotype was due to the co-suppression of *GFP-SMC3* and *AtSMC3*, we named the lines: late suppressor of SMC3 (*lss*), the line whose particular phenotype was visible after 4 weeks and which showed reduced fertility (Figs 2a, S1b); and early suppressor of SMC3 (*ess*), the three lines whose phenotype emerged after 2 weeks and which were sterile in the homozygous state (Figs 2a, S1b). Interestingly, the morphological phenotype of heterozygous *ess* plants was indistinguishable from that of homozygous *lss* plants, supporting the idea that *lss* and *ess* phenotype was attributable to the transgene (Fig. S1b).

**Fig. 2.**
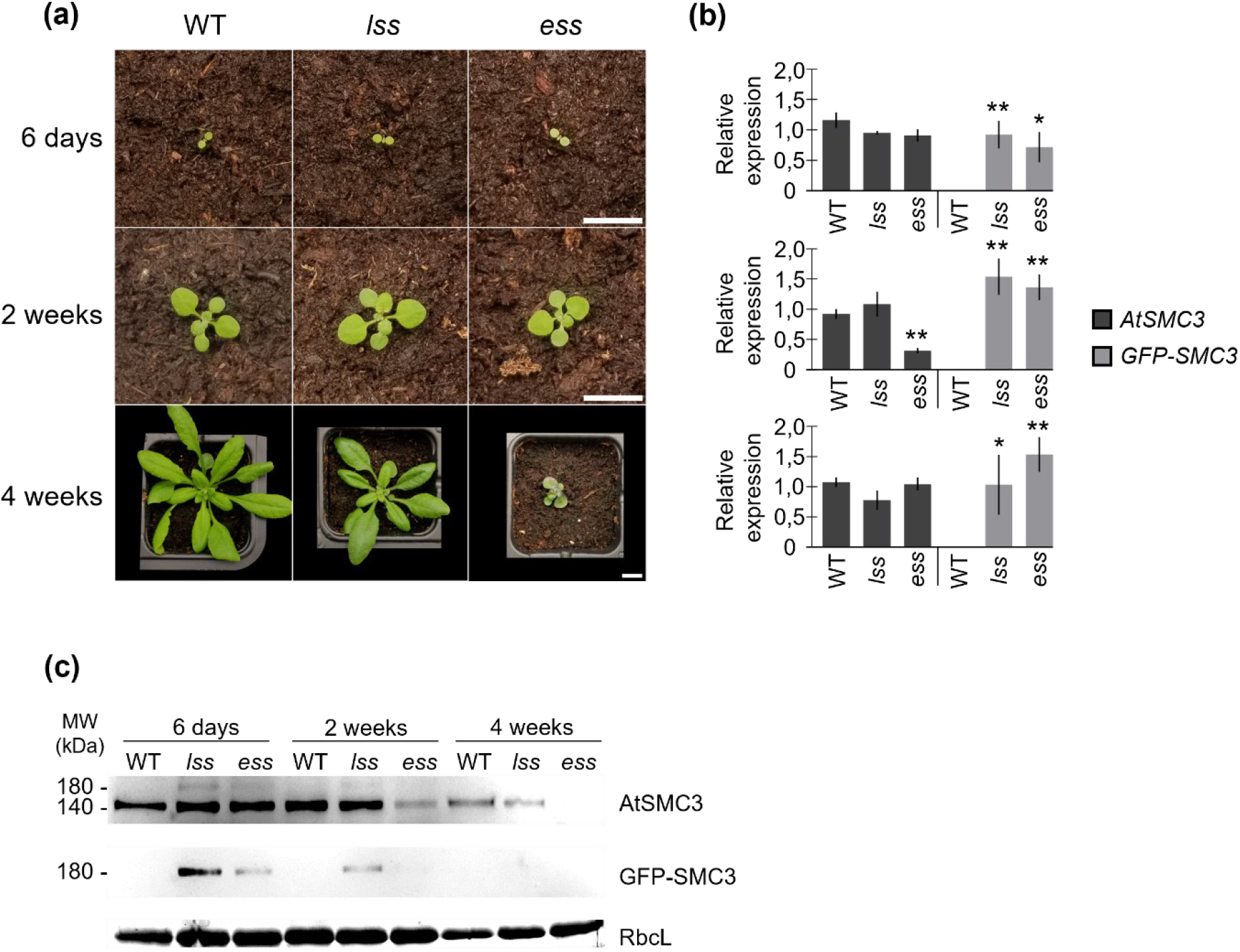
Late Suppressor of SMC3 (*lss*) and Early Suppressor of SMC3 (*ess*) show co-silencing of GFP-SMC3 and AtSMC3. (a) Photographs of long day grown WT, homozygous *lss* and *ess* at 6 days, 2 weeks and 4 weeks. Scale bar, 1 cm. (b) RT-qPCR analysis of *AtSMC3* and *GFP-SMC3* transcript abundance at 6 days, 2 weeks and 4 weeks in WT, *lss* and *ess* genotypes. A segregating population of *ess* was analysed at 6 days. Bars indicate mean and lines indicate the standard error (n = 3 independent biological replicates). Statistical tests (pairwise t-test) shown against respective WT controls, * *P<0*.*05, ** P<0*.*01*. (c) Representative immunoblot with specific antibodies against AtSMC3 and GFP-SMC3 performed on total protein extracts of the indicated genotypes at 6 days, 2 weeks and 4 weeks. A segregating population of *ess* was analysed at 6 days. Rubisco (RbcL) revealed by total protein staining was used as loading control.

### The developmental phenotype of *35S::GFP-SMC3* lines is associated with co-suppression of *GFP-SMC3* and *AtSMC3*

To test whether *lss* and *ess* lines were indeed co-suppression lines, we first quantified *GFP-SMC3* and *AtSMC3* transcript levels by RT-qPCR at different developmental stages in *lss* and one of the three *ess* lines (Fig. 2b). Similar levels of *GFP-SMC3* transcripts were detected in 6-day-old seedlings, 2-week-old plants, and rosette leaves of 4-week-old plants. *AtSMC3* transcripts were also detected in all three tested conditions. The only notable decrease of *AtSMC3* expression was transiently detected in the *ess* line at 2 weeks but recovered to WT level in 4-week-old rosette leaves. Although this transient drop in *AtSMC3* transcript level is significant, it cannot explain the persistent phenotype observed in *ess* at 4 weeks. Moreover, the relatively constant level of *AtSMC3* transcripts in *lss* was not consistent with the hypothesis of a transcriptional or post-transcriptional co-silencing. Therefore, to test whether the putative co-silencing event could take place at a translational or post-translational level, we raised an antibody against the C-terminus region of AtSMC3. In accordance with the theoretical size of AtSMC3 (The Arabidopsis Genome Initiative, 2000; Lam, Yang and Makaroff, 2005), immunobloting using the anti-AtSMC3 antibody detected a 140 kDa protein with similar abundance in each lines at 6 days (Fig. 2c). At 2 weeks, the signal was similar in WT and *lss* lines but strongly reduced in the *ess* line. The decrease in AtSMC3 levels is therefore correlated with the early appearance of a morphological phenotype in *ess* (Fig. 2a,c). At 4 weeks, the AtSMC3 signal also decreased in the *lss* line, which again is correlated with the later appearance of the morphological phenotype in this line. Immunoblots using anti-GFP antibody revealed that the GFP-SMC3 protein was already undetectable at 2 weeks in *ess* and became undetectable at 4 weeks in the *lss* line (Fig. 2c). These results confirm that expression of *GFP-SMC3* in the *lss* and *ess* lines triggers stable translational or post translational co-silencing of AtSMC3 and GFP-SMC3. Furthermore, we confirmed that silencing of AtSMC3 is triggered earlier in *ess*, leading to a severe morphological phenotype that is essentially characterized by growth arrest and sterility (Fig. 2a).

### Transcriptomic analysis of *ess* and *caa39* reveals a common upregulation of oxidative stress response genes

To date, the only reported *AtSMC3* mutant lines contained T-DNA insertions in exons that lead to embryo lethality in the homozygous state (Liu *et al*., 2002), and almost no decrease in *AtSMC3* expression and a WT-like morphological phenotype in the heterozygous state (Schubert *et al*., 2009). These insertion mutants are therefore not convenient for genetic analysis of *AtSMC3* in plants. The *lss* and *ess* lines where AtSMC3 decreases during plant development appear to be more suitable tools to study the consequences of reduced accumulation of AtSMC3 in plants. We therefore decided to compare the impact of AtSMC3 silencing and a hypomorphic Topo VI mutation on gene expression by RNA-seq analysis of *ess* and *caa39* lines at 2 weeks. At this developmental stage, *ess* displays reduced AtSMC3 protein levels, undetectable GFP-SMC3 levels, and only slight morphological defects. We decided not to include 4-week-old *ess* plants in this analysis because of the severity of its morphological phenotype.

To identify commonly and differently regulated genes and pathways in *caa39* and *ess*, we performed gene clustering based on gene expression profiles in *caa39, ess* and the WT control. By combining four different methods, seven main gene clusters could be identified, each cluster corresponding to a typical expression pattern (Fig. 3a, S2, Table S2). Clusters 1, 2 and 3 correspond to genes up regulated in both *caa39* and *ess* (Fig. 3a) and contain 36% of the 9034 genes that are differentially regulated in at least one line (Fig. 3b). Conversely, clusters 5 and 7 contain genes down regulated in both lines and represent about 22% of the genes (Fig. 3a, b). Thus, 58% of the genes belong to clusters containing commonly deregulated genes between *caa39* and *ess*, suggesting a consistent overlap between Topo VI and AtSMC3 functions. The remaining 42% of genes are distributed in two clusters, 4 and 6, which correspond to genes specifically repressed in *caa39* or *ess*, respectively.

**Fig. 3.**
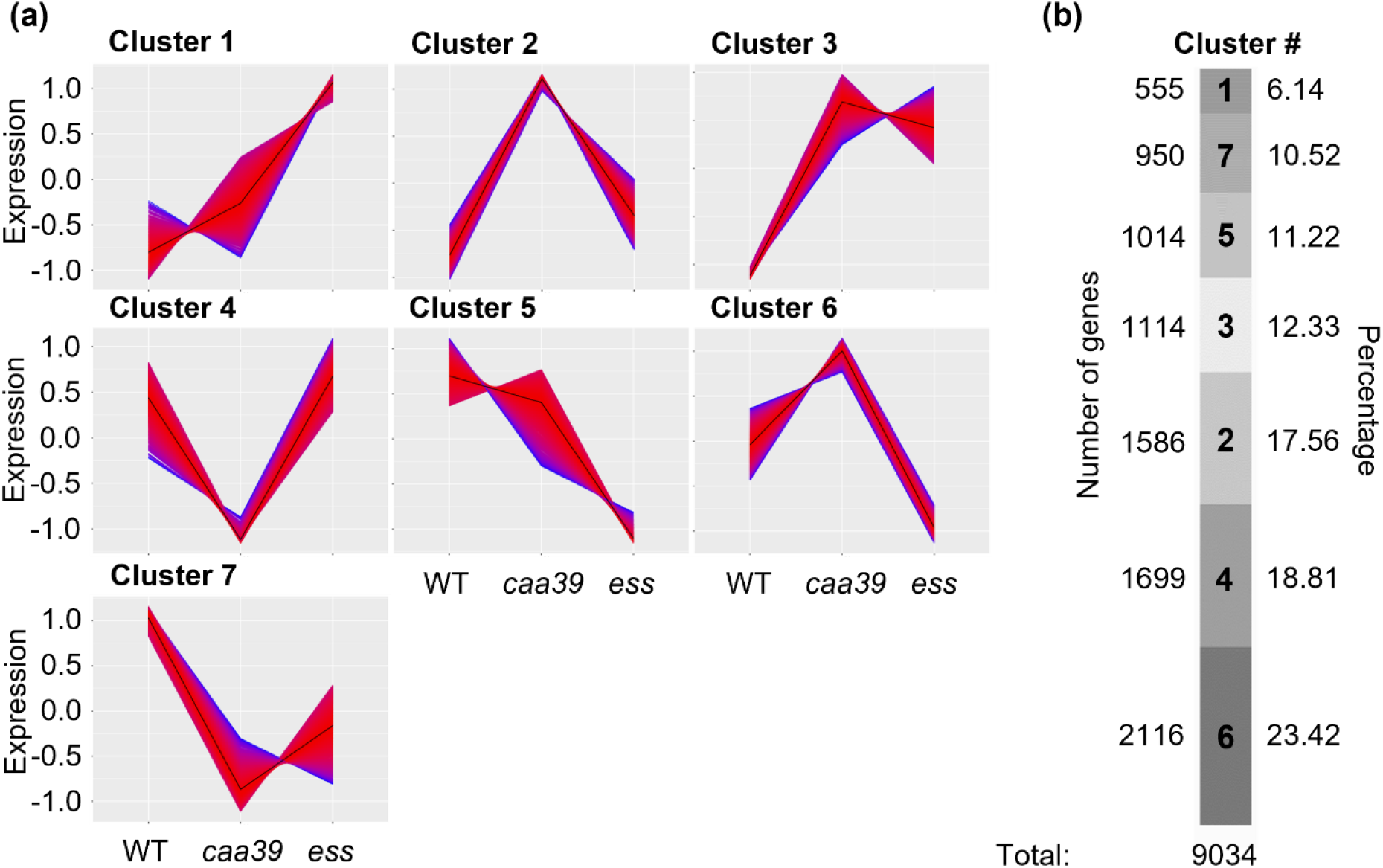
Gene clustering of RNA-seq transcriptomic data for 2-week-old WT, *caa39* and *ess* lines. (a) Cluster 1 corresponds to genes induced in *caa39* and *ess* with different intensities. Cluster 2 corresponds to genes induced in *caa39* and slightly induced in *ess*. Cluster 3 corresponds to genes almost equivalently induced in *caa39* and *ess*. Cluster 4 corresponds to genes only repressed in *caa39*. Cluster 5 corresponds to genes repressed in *ess*. Cluster 6 corresponds to genes upregulated in *caa39* and repressed in *ess*. Finally, cluster 7 corresponds to genes repressed in *caa39* and *ess*. Genes whose expression profile is close to the cluster trend are represented by a red line. Those whose expression profile is more divergent are represented by a blue trace. The exact blue/red scale of each cluster is provided in figure S2. (b) Number and percentage of genes associated with the different clusters. Two biological replicates were performed for each genotype.

We then performed a Gene Ontology (GO) term enrichment analysis to identify biological processes enriched in each cluster. Clusters 5 and 6, which contain genes strongly repressed in *ess* (Fig. 3), are enriched for genes involved in microtubule dynamics, especially in the context of cellular division (Fig. S2). Such down regulation of microtubule related genes could arise from arrest of the cell cycle at an early phase, as has been observed in breast cancer cells where the gene encoding cohesin loader NIPPED-B-LIKE (NIPBL) is silenced (Zhou *et al*., 2017). This hypothesis is further supported by the growth arrest of *ess* plants after 2 weeks (Fig. 2a). Clusters 4 and 7 correspond to genes repressed in *caa39* or in both *caa39* and *ess*, and are enriched for genes involved in several chloroplast functions including light harvesting (Fig. S2, Table S2). Considering that a decrease in photosynthetic efficiency and chloroplast translation is a common response to exposure to diverse stresses (Grennan and Ort, 2007; Upadhyaya and Rao, 2019; Romand *et al*., 2022), the downregulation of genes for chloroplast function could result from a global misregulation of stress signalling in *caa39* and *ess*. Supporting this idea, clusters 1, 2 and 3, which contain genes upregulated in both *caa39* and *ess*, are strongly enriched for genes involved in various stress responses, such as hypoxia, iron starvation or pathogenesis (Fig. S2). This suggests that Topo VI and SMC3 could participate in the regulation of several metabolic or signalling pathways related to stress responses. Importantly, cluster 1 is also specifically enriched for a GO term related to oxidative stress (GO:0006979) (Fig. S2), a feature characteristic of most environmental stresses (Apel and Hirt, 2004). The GO term “Response to oxidative stress” belongs to a more general class called “Response to stress” (GO:0006950) and is subdivided into more specific subclasses among which “response to reactive oxygen species” contains the largest number of genes (GO:0000302) (Fig. 4a).

**Fig. 4.**
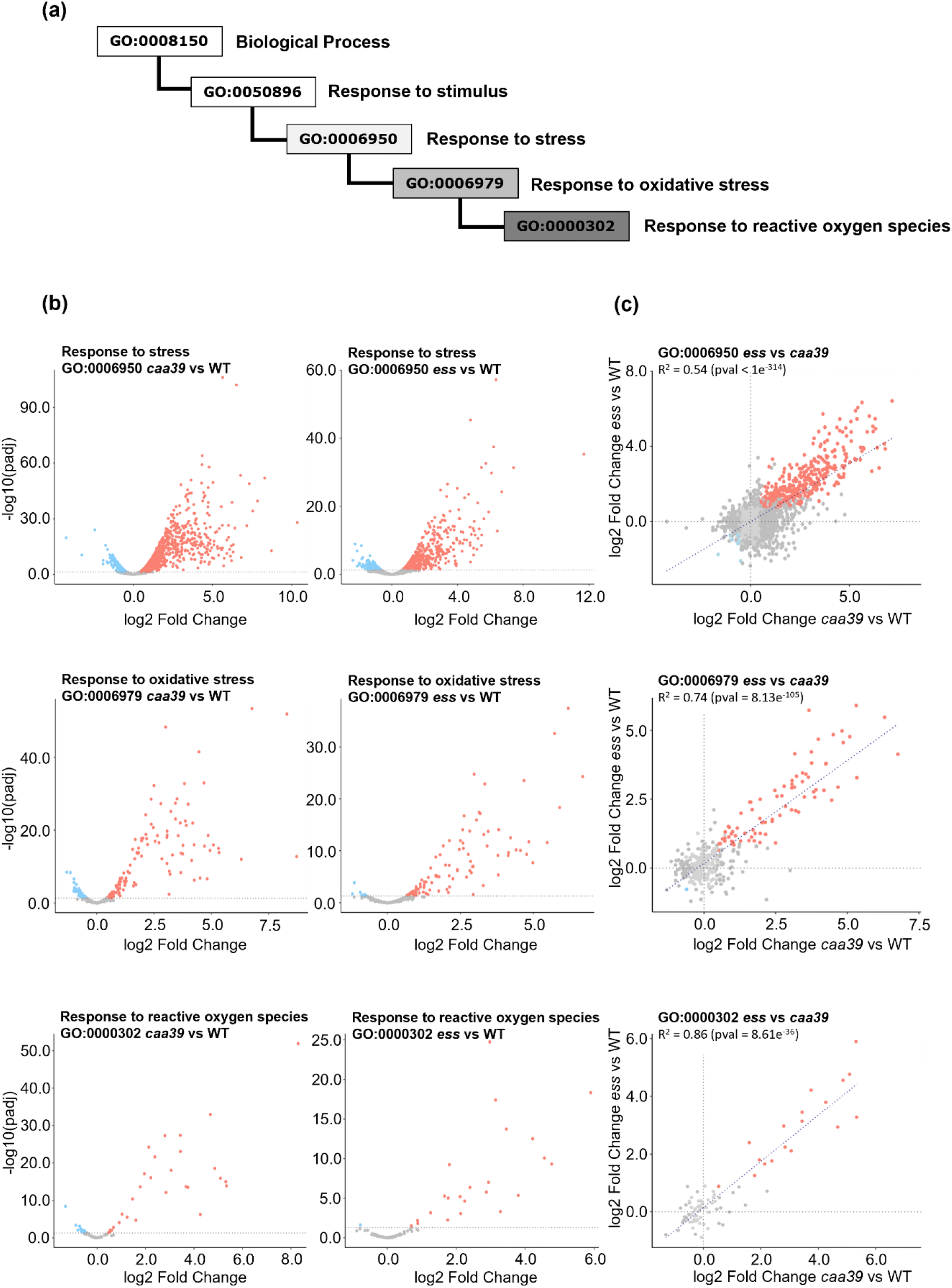
ROS responsive genes are similarly activated in *caa39* and *ess* under normal growth conditions. (a) Diagram representing the hierarchy of the 5 GO: « Response to reactive oxygen species » (GO:0000302) is part of « Response to oxidative stress » (GO:0006979), which is part of « Response to stress » (GO:0006950), which is part of « Response to stimulus » (GO:0050896), which is part of « Biological process » (GO:0008150). (b) Expression profiles of the genes belonging to GO:0006950, GO:0006979 and GO:0000302 in *caa39* (left panel) and *ess* (right panel) compared to the WT at 2 weeks. The p-value is calculated using an exact test for negative binomial distribution and corrected based on Benjamini-Hochberg test. Red dots indicate genes significantly up-regulated, blue dots indicate genes significantly down-regulated, grey dots indicate genes whose expression is not significantly altered. (c) Scatterplots of fold changes versus WT of genes belonging to GO:0006950, GO:0006979 and GO:0000302 for *caa39* and *ess*. Line of best fit and R^2^ with corresponding p-value are indicated. Red dots indicate genes similarly up-regulated, blue dots indicate genes similarly down regulated, grey dots indicate genes differently regulated or genes whose expression is not significantly altered. Two biological replicates were performed for each genotype.

We then performed the reciprocal analysis to determine if an oxidative stress response is activated in *caa39* and *ess*, by analysing the expression of all genes assigned to each of these three GO terms in *caa39* and *ess*. Genes assigned to GO “Response to stress” and more specifically “Response to oxidative stress” and “Response to reactive oxygen species” were in the vast majority activated in both mutant lines compared to WT (Fig. 4b, left and right panels). To further determine whether these genes are commonly regulated in *caa39* and *ess*, we directly compared their fold changes in these two lines. The expression of genes belonging to general GO term “stress response” appeared to be moderately correlated between *caa39* and *ess* (R^2^=0.54) (Fig. 4c). However, the correlation between the two lines increased when considering only the subclass “response to oxidative stress” (R^2^=0.74) and the most specific subclass “response to reactive oxygen species” (R^2^=0.86) (Fig. 4c), suggesting that most oxidative stress related genes are commonly regulated by Topo VI and SMC3.

### *caa39* and SMC3 co-suppression lines show constitutive activation of ^1^O_2_-responsive genes under normal light conditions

To test whether induction of oxidative stress response genes could be attributed to SMC3 co-silencing, we measured the expression of several ^1^O_2_-responsive marker genes (*BON ASSOCIATION PROTEIN 1* / *AT3G61190, BAP1*; *ETHYLENE RESPONSIVE ELEMENT BINDING FACTOR 5* / *AT5G47230, ERF5*; *LIPOXYGENASE 3* / *AT1G17420, LOX3*; *AAA-ATPase* / *AT3G28580, AAA*) and H_2_O_2_-responsive genes (*ASCORBATE PEROXIDASE 1* / *AT1G07890, APX1*; *FERRETIN 1* / *AT5G01600, FER1*; *AT3G49160, pKsi*) (Op Den Camp *et al*., 2003; Šimková *et al*., 2012) by RT-qPCR at 6 days and 2 weeks in *lss, ess* and *caa39*. In previous studies, 6-day-old *caa39* plants were reported to show selective activation of a set of ^1^O_2_-responsive genes (Šimková *et al*., 2012). As expected, 6-day-old *caa39* seedlings strongly accumulated transcripts for the ^1^O_2_-responsive genes *BAP1* and *AAA*, whereas they displayed very slight variations in the expression of H_2_O_2_-responsive genes (Fig. S3). However, *lss* and *ess* lines showed transcript levels similar to the WT control for both H_2_O_2_ and ^1^O_2_ markers, suggesting that over-expression of *GFP-SMC3* at this stage does not trigger oxidative stress response under normal light conditions (Fig. S3). In agreement with the RNA-seq data, 2-week-old *caa39* plants displayed significant increased transcript levels of ^1^O_2_ marker genes. Similarly, 2-week-old *ess* line accumulated high amount of all tested ^1^O_2_ marker transcripts, up to over 250-fold activation for *AAA* expression. Interestingly, *lss* line only partially activated the ^1^O_2_ markers with a slight induction of *AAA* gene (7.3 fold change) and *ERF5* gene (1.73 fold change) (Fig. 5). No significant increase in the expression of H_2_O_2_-responsive genes was observed, suggesting a specific and constitutive activation of the ^1^O_2_ pathway in AtSMC3 suppressors and *caa39*.

**Fig. 5.**
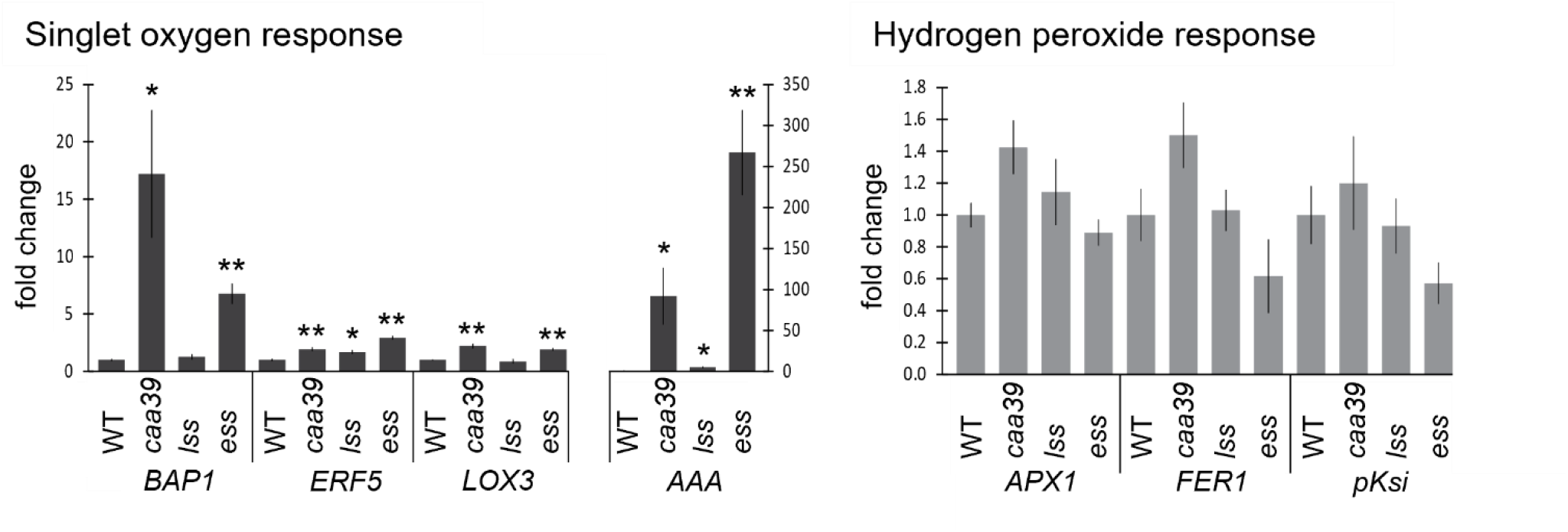
The suppressors of SMC3 and *caa39* show activation of the ^1^O_2_ response. RT-qPCR analysis measuring the transcript abundance of the ^1^O_2_-responsive genes *BAP1, ERF5, LOX3, AAA* and the H_2_O_2_-responsive genes *APX1, FER1, pKsi* at 2 weeks of growth. Bars indicate mean and lines indicate the standard error (n = 3 independent biological replicates). Statistical tests (pairwise t-test for data following a parametric distribution; Games-Howell Post-hoc test for data not following a parametric distribution) shown against respective WT controls, * *P<0*.*05, ** P<0,01*.

### SMC3 co-suppression and *caa39* lines display enhanced resistance to high light stress

To further investigate the interplay between Topo VI and AtSMC3 in the constitutive activation of the ^1^O_2_ response, we crossed *lss* and *ess* plants with the *caa39* mutant to obtain the *caa39 lss* and *caa39 ess* lines. The *caa39* mutation coupled to SMC3 suppression led to an additive phenotype combining the morphological characteristics of each parental lines, leading to severe growth defects (Fig. S4). To investigate the physiological response to the production of ^1^O_2_ we exposed the SMC3 suppressors, the *caa39* mutant, and the double *caa39 lss* and *caa39 ess* lines to high light photooxidative stress conditions (1500 μ mol photons. m^-2^ .s^-1^, 4°C, 48 h, 8 h photoperiod, Fig. 6a) that are specifically designed to generate a burst of ^1^O_2_ production (Ramel *et al*., 2012). We performed the high light stress on 2-week-old *ess* and 4-week-old *lss* in order to limit indirect effects due to morphological differences or residual expression of GFP-SMC3.

**Fig. 6.**
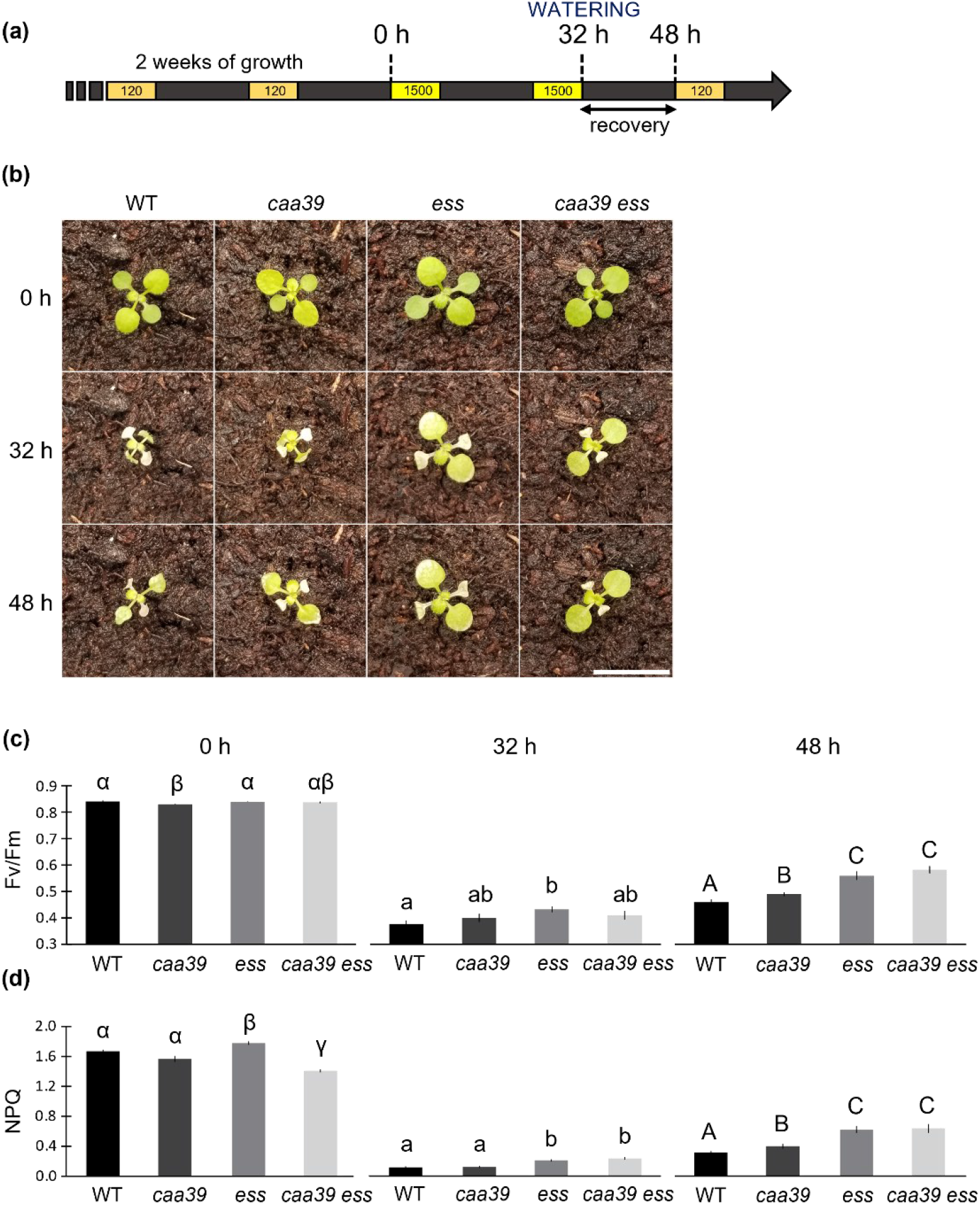
SMC3 suppressors show enhanced tolerance to photooxidative stress. (a) Timescale of the high light stress experiment performed on two-week-old plantlets. Orange, 8 h normal light periods (120 μmol m^-2^ s^-1^); yellow, 8 h high light periods (1500 μmol m^-2^ s^-1^); black, 16 h dark periods. 0 h, onset of the high light stress; 32 h, end of the second high light illumination period; 48 h, end of the dark period following the high light stress. (b) Pictures of representative 2-week-old WT, *caa39, ess* and *caa39 ess* plantlets at 0 h, steady state; 32 h, after high-light exposure; 48 h, following a 16 h recovery period in the dark. Scale bar, 1 cm. (c) PSII efficiency (Fv/Fm) of WT, *caa39, ess* and *caa39 ess* at 0 h, 32 h and 48 h. Black lines indicate the standard error (n = 12 plants from 3 independent biological replicates). Compact letter display represents statistic groups and was generated using a pairwise t-test corrected with the Benjamini/Hochberg FDR method. (d) Non-Photochemical Quenching (NPQ) measurements. Black lines indicate the standard error (n = 12 plants from 3 independent biological replicates). Compact letter display represents statistic groups and was generated using a pairwise t-test corrected with the Benjamini/Hochberg FDR method.

After 32 h high light stress, 2-week-old plantlets of all tested genotypes displayed photo-bleaching on the entire surface of the cotyledons and partly on the first and second leaves (Fig. 6b). Although the WT and *caa39* plantlets appeared to be more severely affected and desiccated, as shown by down-curled leaves after the stress period, the Fv/Fm measurements indicated PSII photoinhibition was similar in all lines (Fig. 6c, middle panel “32 h”). Interestingly, after the recovery period of 16 h in the dark, *caa39* displayed a slightly higher Fv/Fm ratio compared to the WT line (Fig. 6c, right panel “48 h”). Following a similar trend, the SMC3 suppressor lines, *ess* and *caa39 ess* displayed an even greater Fv/Fm ratio indicating that there is a more efficient recovery of PSII capacity in the absence of AtSMC3 (Fig. 6c). To better understand the high light tolerance of SMC3 suppressor, we measured non-photochemical quenching (NPQ) before and after exposure to oxidative stress. Surprisingly, under normal light conditions, NPQ was lower in *caa39 ess* compared to the WT and *caa39*, whereas the *ess* line had the highest level of NPQ (Fig. 6d, left panel “0 h”). However, after exposure to photoinhibitory conditions, the NPQ values of both SMC3 suppressing lines clustered and became substantially higher than WT and *caa39* (Fig. 6d, middle panel “32 h”). Finally, after the 16 h recovery in the dark, NPQ remained higher in SMC3 suppressors than in the other lines, thus correlating the tolerance to photooxidative stress with the activation of NPQ (Fig. 6d, right panel “48 h”). These results show that the silencing of SMC3 and, to a lesser extent, the *caa39* mutation led to a resistance to high light, likely through an overactivation of the NPQ pathways.

To confirm that these physiological observations are a consequence of SMC3 silencing, we repeated this experiment using 4-week-old WT, *caa39, lss* and *caa39 lss* plants. After applying the same stress conditions on 4-week-old plants, we observed that the oldest WT leaves were partly bleached. However, control plants remained mostly green, showing the higher tolerance of 4-week-old plants than 2-week-old plantlets to these photoinhibitory conditions (Fig. S5a). Immediately after exposure to high light, Fv/Fm was slightly but significantly higher in *lss* and *caa39 lss* than in WT and *caa39* (Fig. S5b). In agreement with observations in 2-week-old *caa39 ess*, NPQ under normal light conditions was lower in *caa39 lss* compared to the WT. (Fig. 6d, left panel “0 h”). However, after exposure to photoinhibitory conditions, the NPQ appeared to be not significantly higher in *lss* lines than in WT and *caa39* (Fig. 6d, middle panel “32 h”). These results show that SMC3 suppressors and to a lesser extent *caa39*, are resistant to photoinhibitory conditions.

## DISCUSSION

Photooxidative stress conditions trigger retrograde signalling to activate expression of nuclear stress responsive genes essential to prevent damages to macromolecules and to detoxify the cell (Gill and Tuteja, 2010; Kerchev and Van Breusegem, 2022). The coordinated expression of these genes requires the activity of transcription factors (Dubos *et al*., 2010; Jiang *et al*., 2017; Feng *et al*., 2020), chromatin remodelling factors (Song, Liu and Han, 2021) and topoisomerases (Šimková *et al*., 2012). However, interactions between these different factors remain unclear. In this study, we presented two SMC3 silencing lines, *lss* and *ess*, where the ^1^O_2_-responsive gene *AAA* is constitutively activated under normal light conditions. This result echoes a previous study showing that *caa39*, a hypomorphic mutant of Topo VI, constitutively activates several ^1^O_2_-responsive genes including *AAA* (Baruah *et al*., 2009; Šimková *et al*., 2012). Additionally, RNA-seq analysis revealed that genes overexpressed in both *caa39* and *ess* are enriched for oxidative stress responsive genes. The confirmation of RNA-seq data by RT-qPCR revealed that many ^1^O_2_-responsive genes, like *BAP1, ERF5* or *LOX3* (Op Den Camp *et al*., 2003; Danon *et al*., 2005) are upregulated in *caa39* and *ess*. The similarities of the transcriptomic features between *caa39* and SMC3 suppressors raises the idea that Topo VI and the cohesin complex could be involved in a common pathway regulating the response to ^1^O_2_.

Supporting this idea, we demonstrated that the BIN4 subunit of Topo VI directly interacts with SMC3(193-807). SMC3(193-807) lacks the N and C-terminal extremities which assemble in the ATP binding cassette of SMC3. This domain contains the interaction site with the α-kleisin subunit SCC1 which participates in the formation of the cohesin ring (Gligoris *et al*., 2014). Moreover, the enclosure and opening of the SMC3/SCC1 interface require the binding and hydrolysis of ATP (Marcos-Alcalde *et al*., 2017; Muir *et al*., 2020), a process which cannot be achieved in SMC3(193-807). This suggests that the interaction between SMC3(193-807) and BIN4 can take place without the integration of SMC3 into a functional cohesin complex. In regard of the complete loss of BiFC signal obtained with the pair BIN4-cYFP associated with SMC3(193-807, Δ525-632)-nYFP lacking the hinge domain of SMC3, we also propose that BIN4 interacts with the hinge domain of SMC3. Indeed, several studies have shown that this domain is prone to interact with partner proteins, as is the case for SMC1 (Chiu, Revenkova and Jessberger, 2004) and for hinderin (Patel and Ghiselli, 2005).

At a physiological level, 2-week-old *ess* and *caa39 ess* plants displayed an enhanced recovery of PSII efficiency, accompanied by activation of NPQ, after recovery from severe photoinhibitory conditions. Moreover, 4-week-old *lss* and *caa39 lss* displayed reduced PSII photoinhibition after high-light treatment compared to WT plants. The enhanced resistance of SMC3 suppressors could be attributed to lower light-harvesting efficiency or could arise from a constitutive stress state that might prime the plants for further stress exposure. However, under normal light conditions, the similar PSII efficiency of WT, *lss* and *ess* shows that SMC3 suppressors are not pre-stressed and have no defect in light harvesting. We therefore propose that SMC3 is required for the regulation of the photooxidative stress response in *Arabidopsis thaliana*. Unlike 6-day-old *caa39* seedlings (Baruah *et al*., 2009; Šimková *et al*., 2012), 2-week and 4-week-old *caa39* plants did not show pronounced tolerance to photooxidative stress under our conditions. This suggests that *caa39* is sensitive to variations in experimental setup or developmental stages.

Limited signs of photodamage after high light treatment in *ess*, combined with the spontaneous appearance of necrotic spots very similar to hypersensitive response on 5-week-old-leaves, suggest that SMC3 could participate in the regulation of a wider range of stress responses than Topo VI. Several studies support this idea. For instance, in Cornelia de Lange Syndrome, cell lines carrying mutations in *SMC1A* or *SMC3* exhibit higher level of protein carbonylation, reflecting increased global oxidative stress (Gimigliano *et al*., 2012). In budding yeast, temperature sensitive mutants of the cohesin complex show induction of the cell wall stress responsive genes CHITIN SYNTHASE 3 (*CHS3*) and the β-1,3-GLUCAN SYNTHASE (*FKS2*), display increased amount of chitin in their cell wall, and are sensitive to cell wall stress-inducing agents (Kothiwal, Gopinath and Laloraya, 2021). Finally, in embryonic stem cells, the cohesin complex was reported to participate in the establishment of the heat stress response by modifying the local chromatin architecture around heat stress-activated enhancers (Lyu, Rowley and Corces, 2018).

From a technical perspective, this study generated new and efficient tools that mimic conditional knockout or knockdown lines and are useful for studying the function of AtSMC3 in plants. Indeed, the SMC3 silencing lines *lss* and *ess* drive a robust translational or post-translational silencing of AtSMC3 and can be crossed with other mutant lines while conserving their ability to silence AtSMC3. Furthermore, the overexpression of GFP-SMC3 in 6-day-old *lss* and *ess* and the new anti-AtSMC3 antibody open the field for biochemical studies of AtSMC3.

## ACKNOWLEDGEMENTS

We want to express our gratitude to students who contributed to this work, especially Chloé Carassus. This work was supported by the French National Research Agency (ANR-14-CE02-0010 to CL). F.V. is a recipient of a PhD fellowship from the French Ministry of Higher Education, Research and Innovation.

## AUTHOR CONTRIBUTIONS

FV, DA, C Laloi performed the experiments. C Lecampion performed the bioinformatic analyses. FV, SD, BF and C Laloi analysed the data. NK and BF contributed to material and methods. FV and C Laloi designed the research. FV and C Laloi wrote the manuscript. All authors read and approved the final manuscript.

## DATA AVAILABILITY

RNA-seq data are available at https://www.ncbi.nlm.nih.gov/geo/query/acc.cgi?acc=GSE211131 (reviewer token: wfydqwyqrbgdfir).

## SUPPORTING INFORMATION

**Fig. S1** SMC3 suppressors display growth defects, anthocyanin accumulation and localized cell death throughout the silencing of AtSMC3.

**Fig. S2** Expression profil and GO term enrichment of the 7 gene clusters.

**Fig. S3** 6-day-old *lss* and *ess* are not impaired in the oxidative stress response.

**Fig. S4** Suppressing AtSMC3 and AtTopoVI A leads to additive morphological phenotypes.

**Fig. S5** 4-week-old *lss* displays a slight tolerance to photooxidative stress.

**Table S1** BIN4 interacts with SMC3 in Y2H.

**Table S2** Complete list of genes associated with clusters 1-7.

**Method S1** Script used for the generation of gene clusters and analyses of GO term enrichments.

