## Supplemental Figures 1-5 for "Structural Maintenance of Chromosome 3 interacts with the Topoisomerase VI complex and contributes to the oxidative stress response in *Arabidopsis thaliana*"

The following Supporting Information is available for this article:

**Fig. S1** SMC3 suppressors display growth defects, anthocyanin accumulation and localized cell death throughout the silencing of AtSMC3.

**Fig. S2** Expression profil and GO term enrichment of the 7 gene clusters.

**Fig. S3** 6-day-old *lss* and *ess* are not impaired in the oxidative stress response.

**Fig. S4** Suppressing AtSMC3 and AtTopoVI A leads to additive morphological phenotypes.

**Fig. S5** 4-week-old *lss* displays a slight tolerance to photooxidative stress.

**Table S1** BIN4 interacts with SMC3 in Y2H.

**Table S2** Complete list of genes associated with clusters 1-7.

**Method S1** Script used for the generation of gene clusters and analyses of GO term enrichments.

**Fig. S1** *35S::GFP-SMC3* primary transformants display growth defects, anthocyanin accumulation and localized cell death throughout the silencing of AtSMC3. (a) Pictures of five independent *35S::GFP-SMC3* transformants grown under long days. A recurrent morphological phenotype is observable in the T1 generation. The severity of the phenotype differs among transformants. (b) Pictures of long day grown WT, homozygous *lss,* heterozygous *ess* and homozygous *ess* at 2 weeks, 4 weeks and 5 weeks. The *e10.12* and *e1.5* columns show two other independant transformants of *GFP-SMC3* displaying a phenotype similar to *ess* line. Scale bar, 1cm.

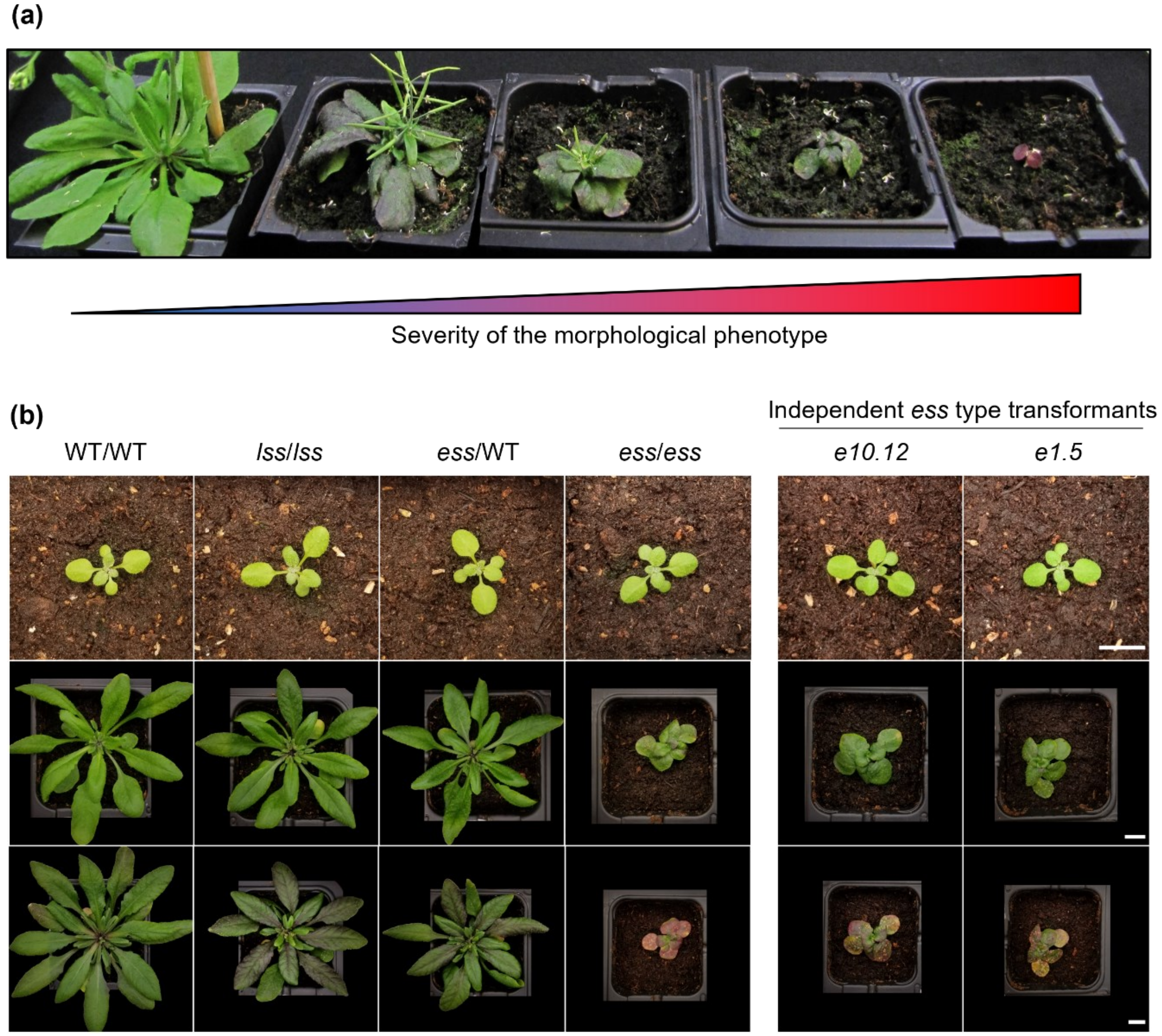

**Fig. S2** Expression profil and GO term enrichment of the 7 gene clusters. For each cluster, Upper panel: black line represents the typical expression profile. Genes whose expression profile is close to the cluster trend are represented by a red line. Those whose expression profile is more divergent are represented by a blue trace. The blue/red scale indicates the gene cluster membership score. A gene is considered to be significantly associated with the cluster when this score is greater than 0.85. The amount and percentage of genes whose gene cluster membership score is higher than 0.85 is indicated in the table. Lower panel: GO term enrichment analysis performed on cluster 1-7. Parent GO terms of a specific GO subclass are shown in the same colour. The size of the cercles represent the number of genes belonging to the GO term. The color of the cercles represents the significance of the enrichment based on the False Discovery Rate (FDR) method. Only the 20 most significant enrichments are shown for each cluster. The position of the cercles on the x-axis indicates the fold enrichment of the corresponding GO term compared to random gene distribution.

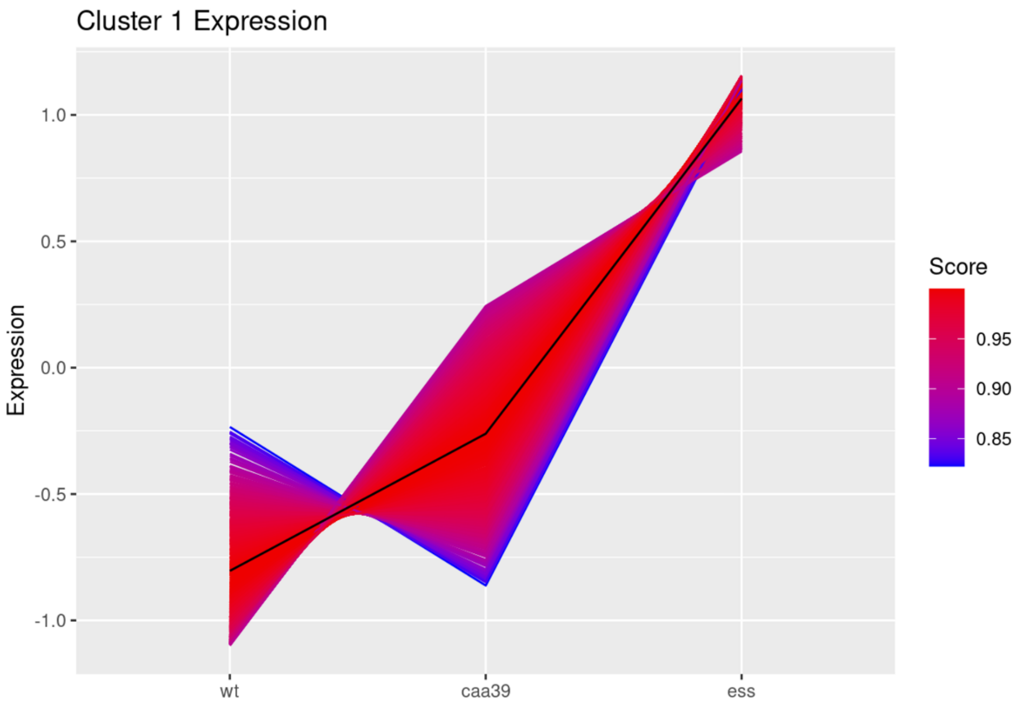


**Percentage of genes retained after cluster filtering using cluster membership score**

| Cluster | Raw | Filtered | Retained (%) |
|---|---|---|---|
| 1 | 564 | 555 | 98.40 |

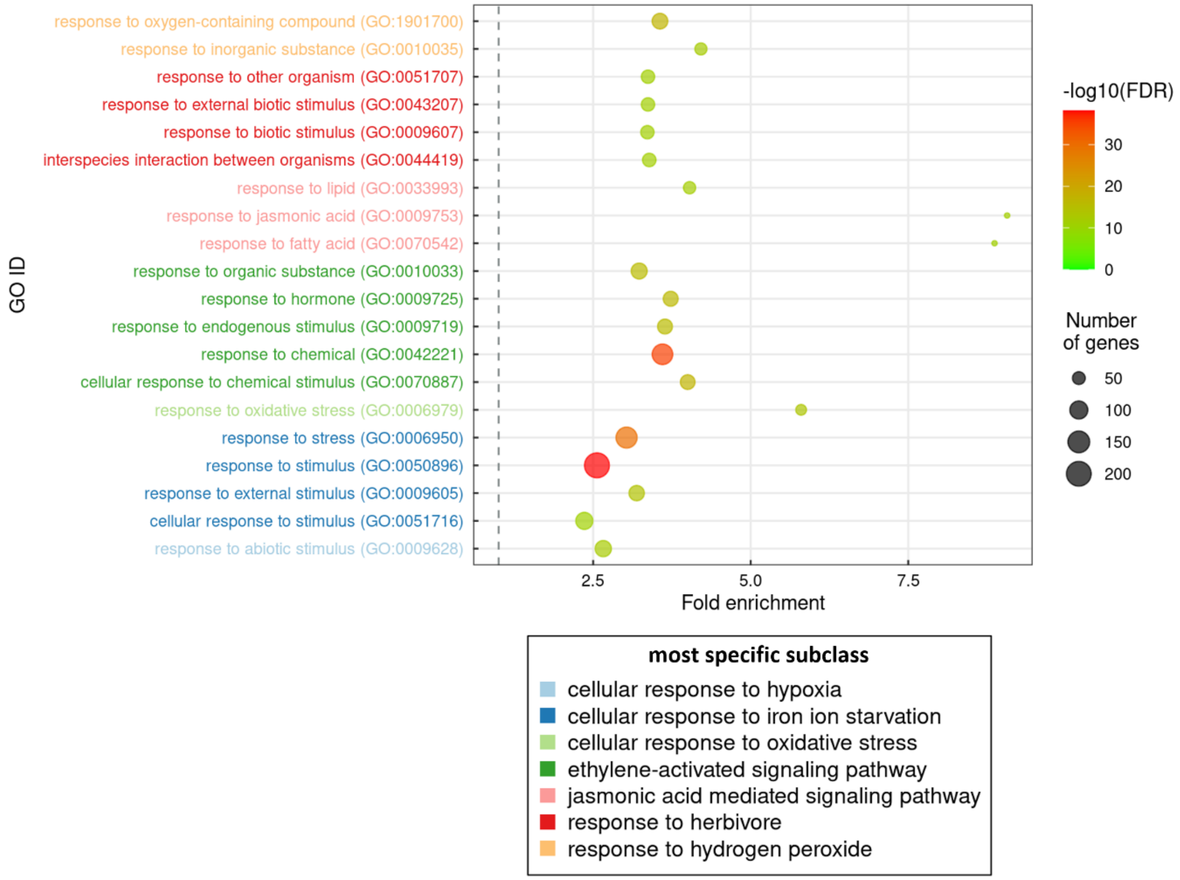

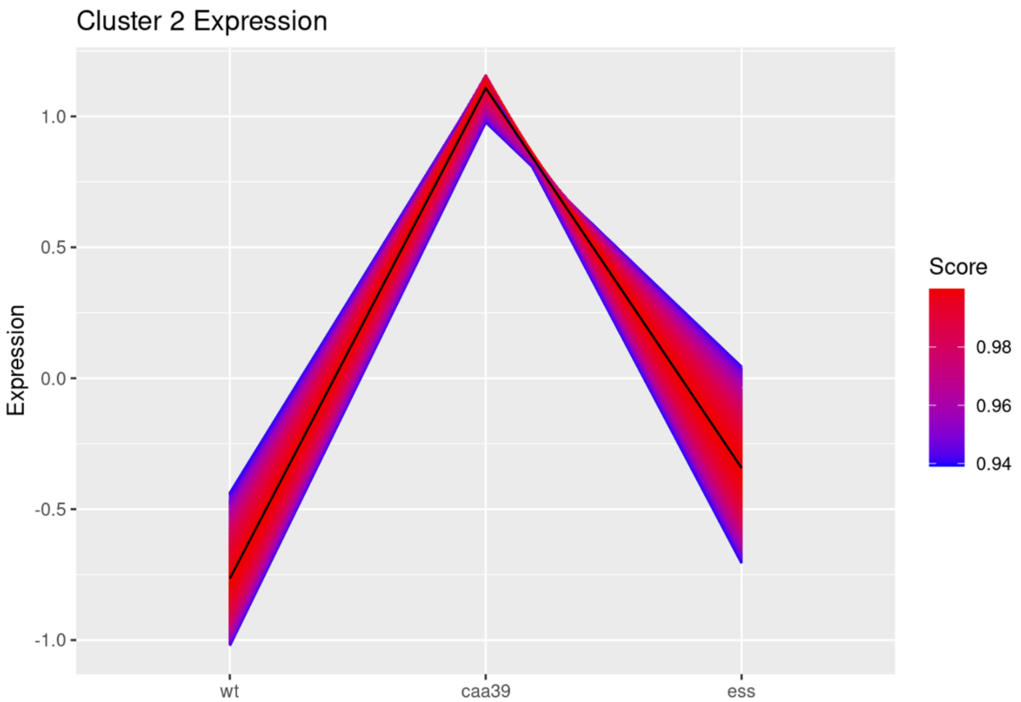


**Percentage of genes retained after cluster filtering using cluster membership score**

| Cluster | Raw | Filtered | Retained (%) |
|---|---|---|---|
| 2 | 1586 | 1586 | 100.00 |

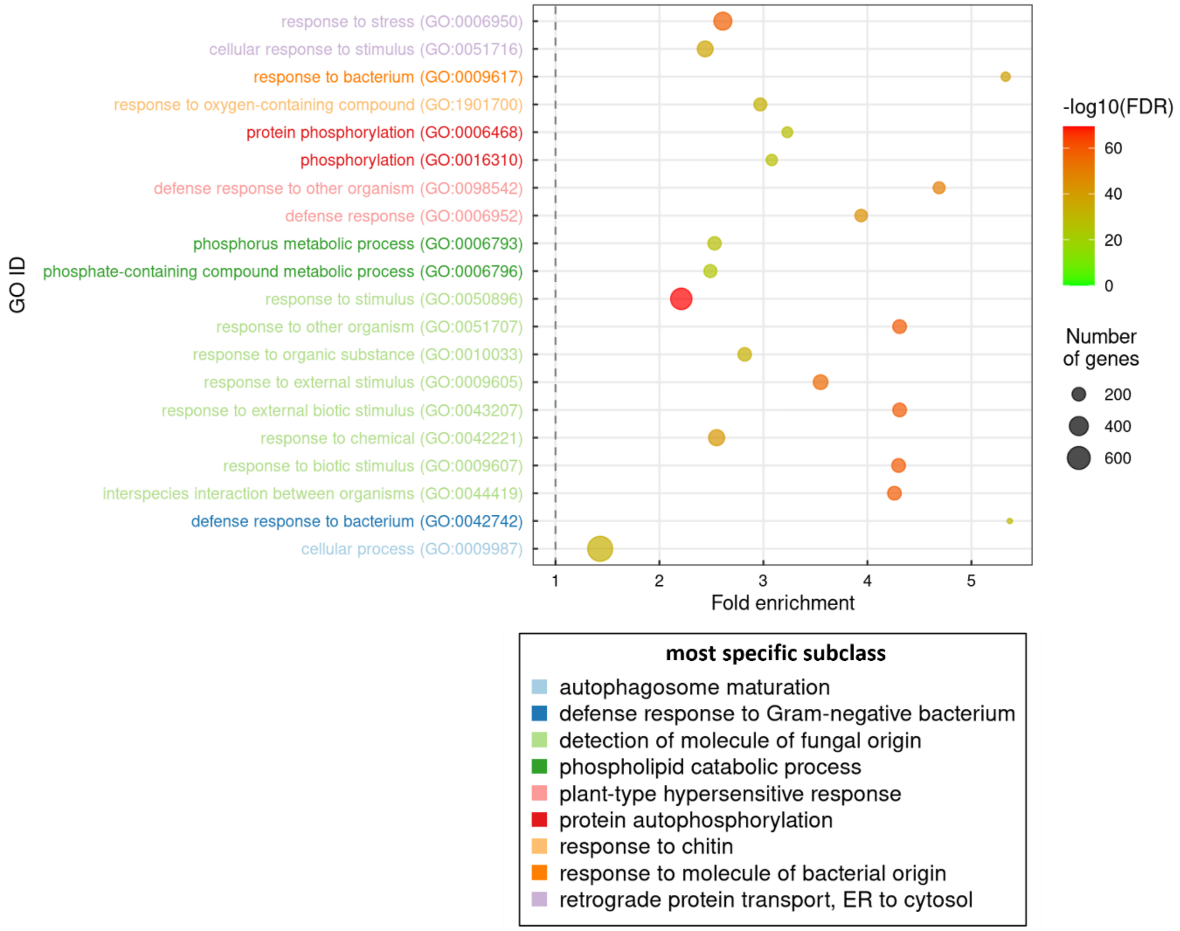

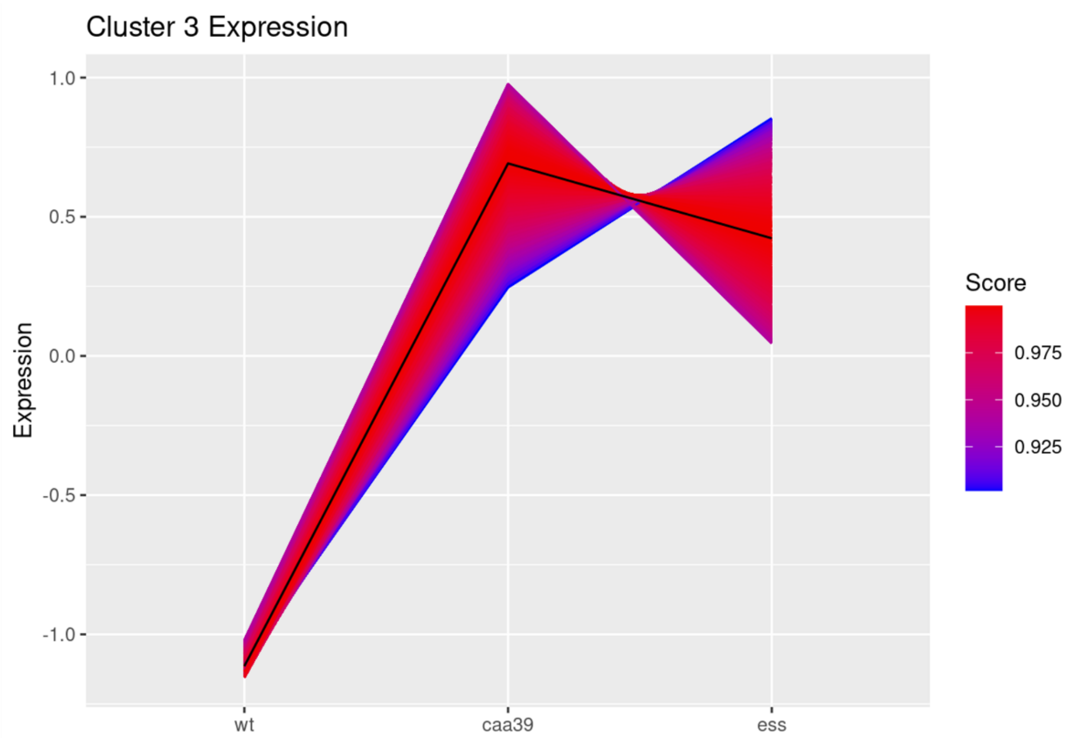


**Percentage of genes retained after cluster filtering using cluster membership score**

| Cluster | Raw | Filtered | Retained (%) |
|---|---|---|---|
| 3 | 1114 | 1114 | 100.00 |

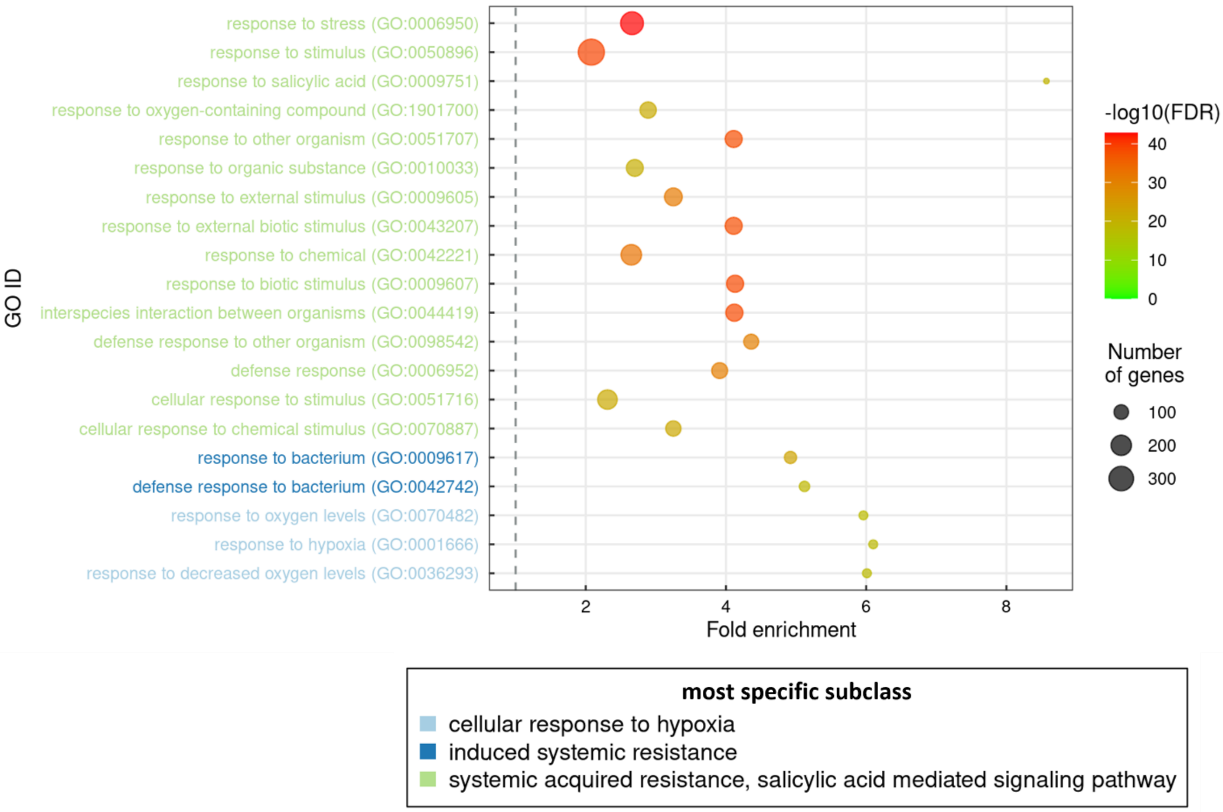

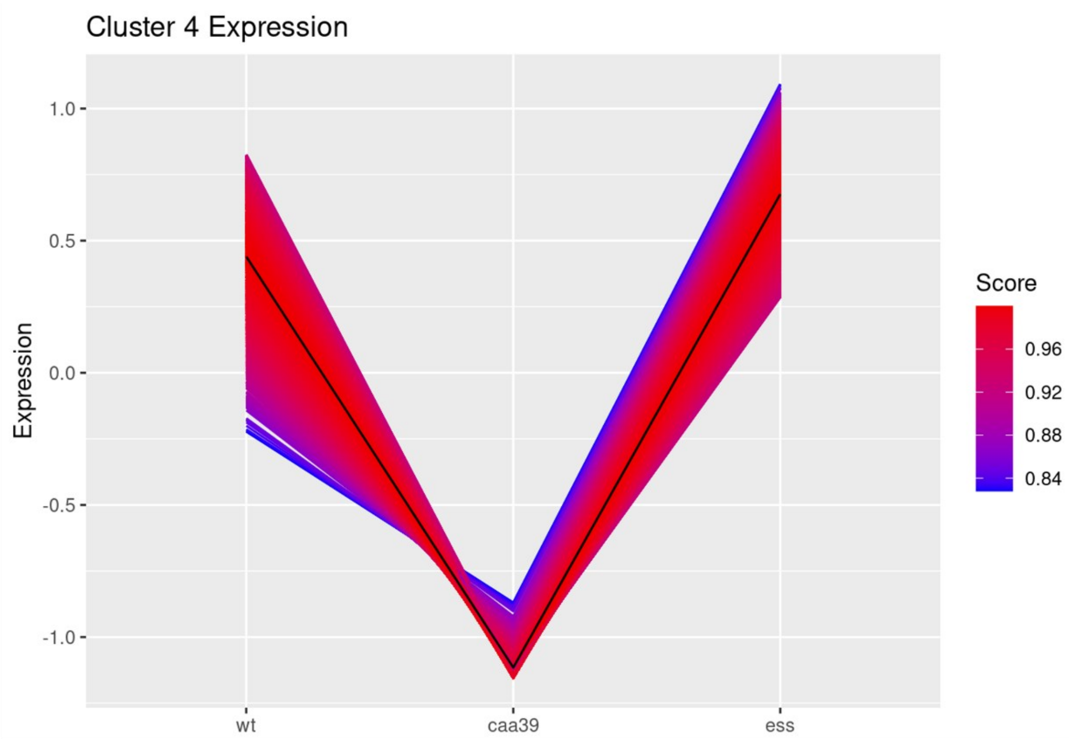


**Percentage of genes retained after cluster filtering using cluster membership score**

| Cluster | Raw | Filtered | Retained (%) |
|---|---|---|---|
| 4 | 1707 | 1699 | 99.53 |

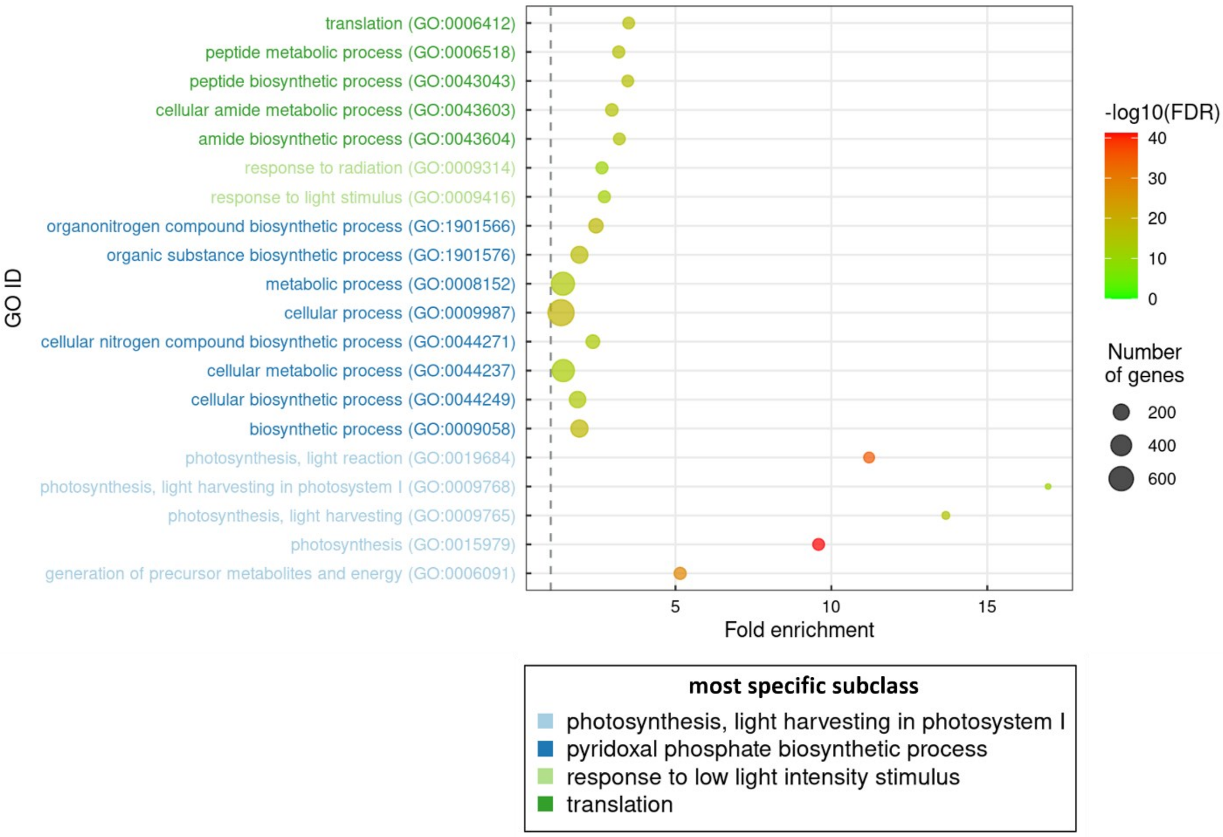

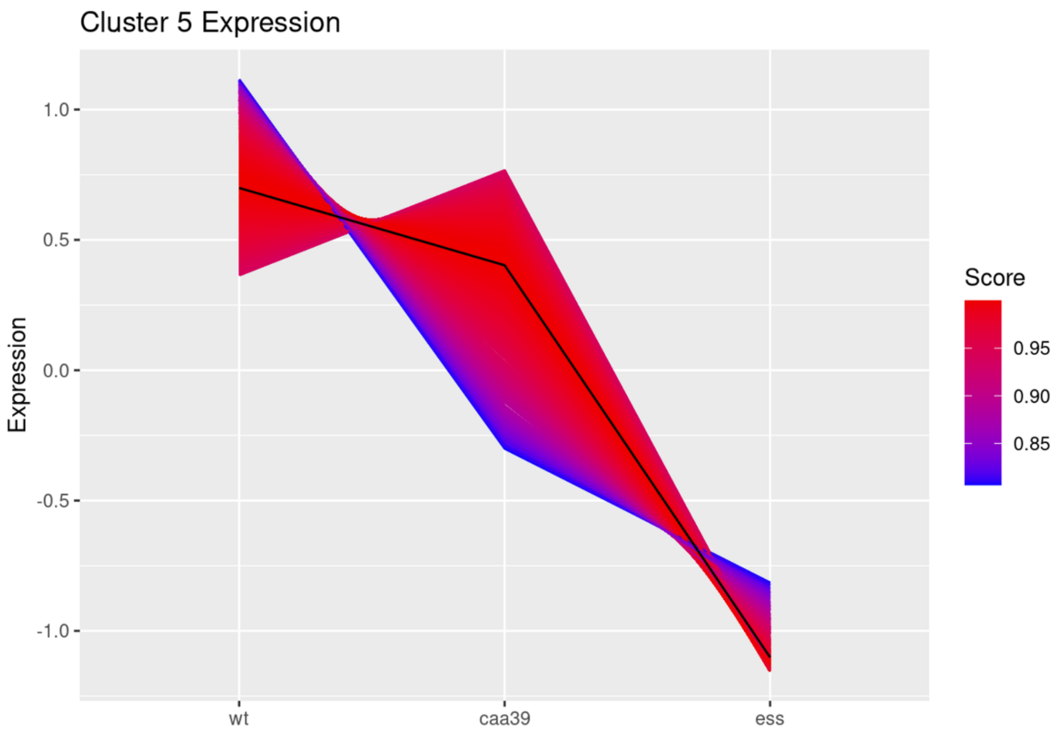


**Percentage of genes retained after cluster filtering using cluster membership score**

| Cluster | Raw | Filtered | Retained (%) |
|---|---|---|---|
| 5 | 1042 | 1014 | 97.31 |

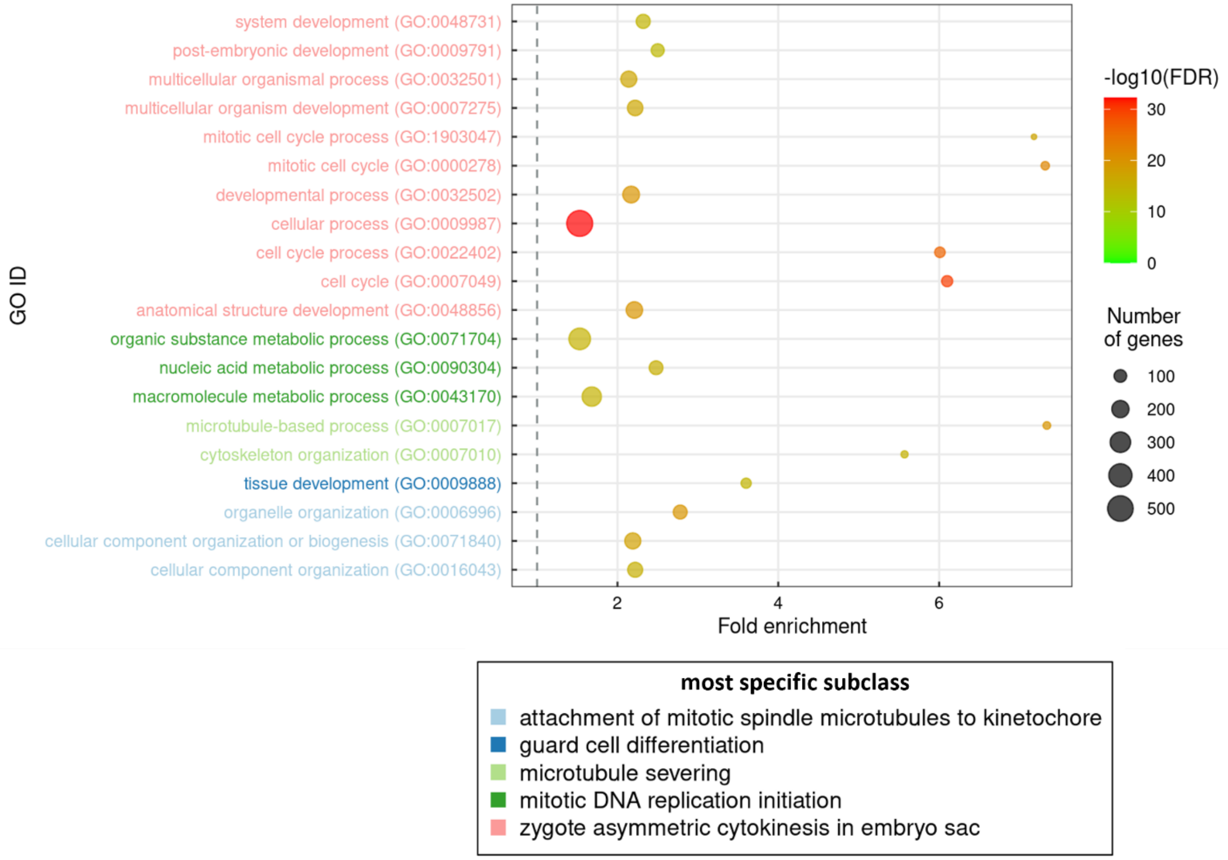

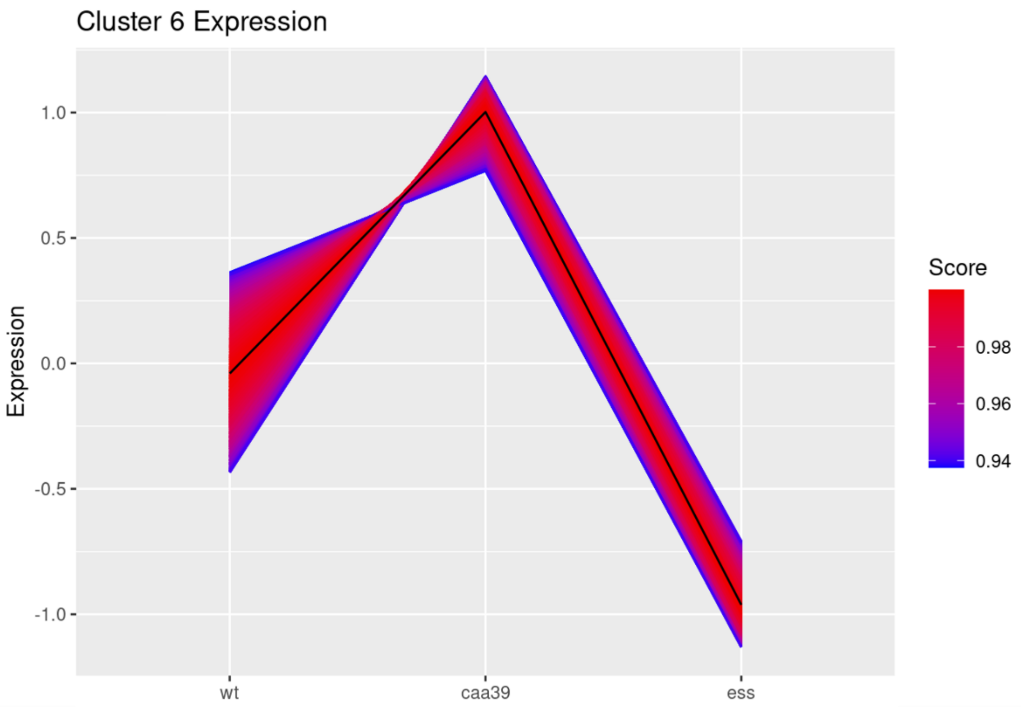


**Percentage of genes retained after cluster filtering using cluster membership score**

| Cluster | Raw | Filtered | Retained (%) |
|---|---|---|---|
| 6 | 2116 | 2116 | 100.00 |

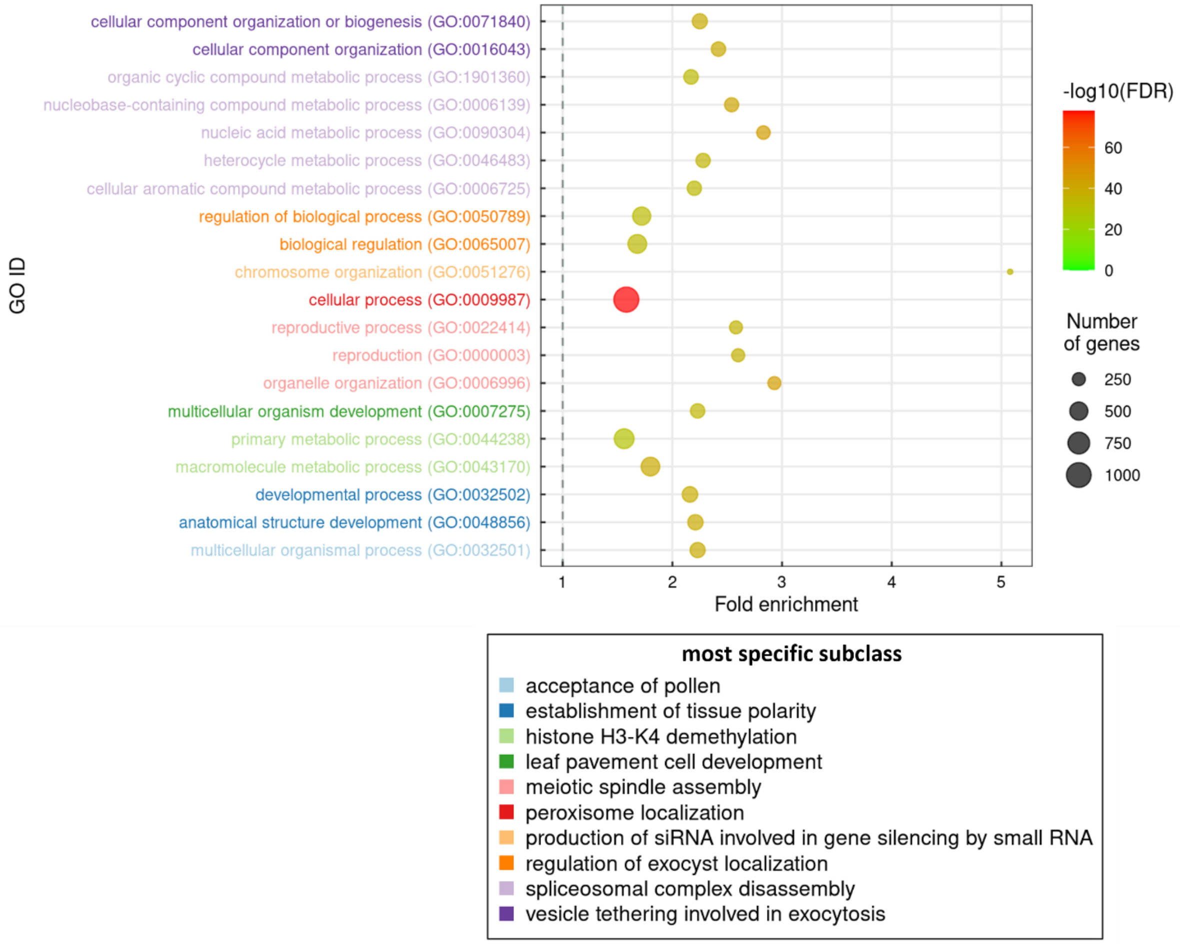

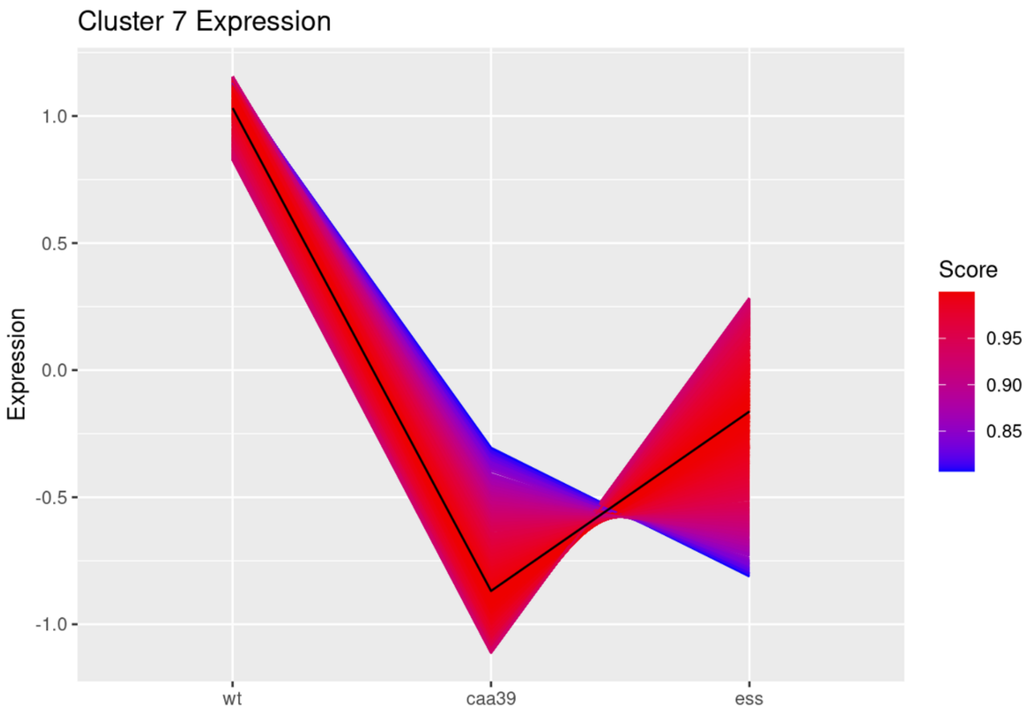


**Percentage of genes retained after cluster filtering using cluster membership score**

| Cluster | Raw | Filtered | Retained (%) |
|---|---|---|---|
| 7 | 990 | 950 | 95.95 |

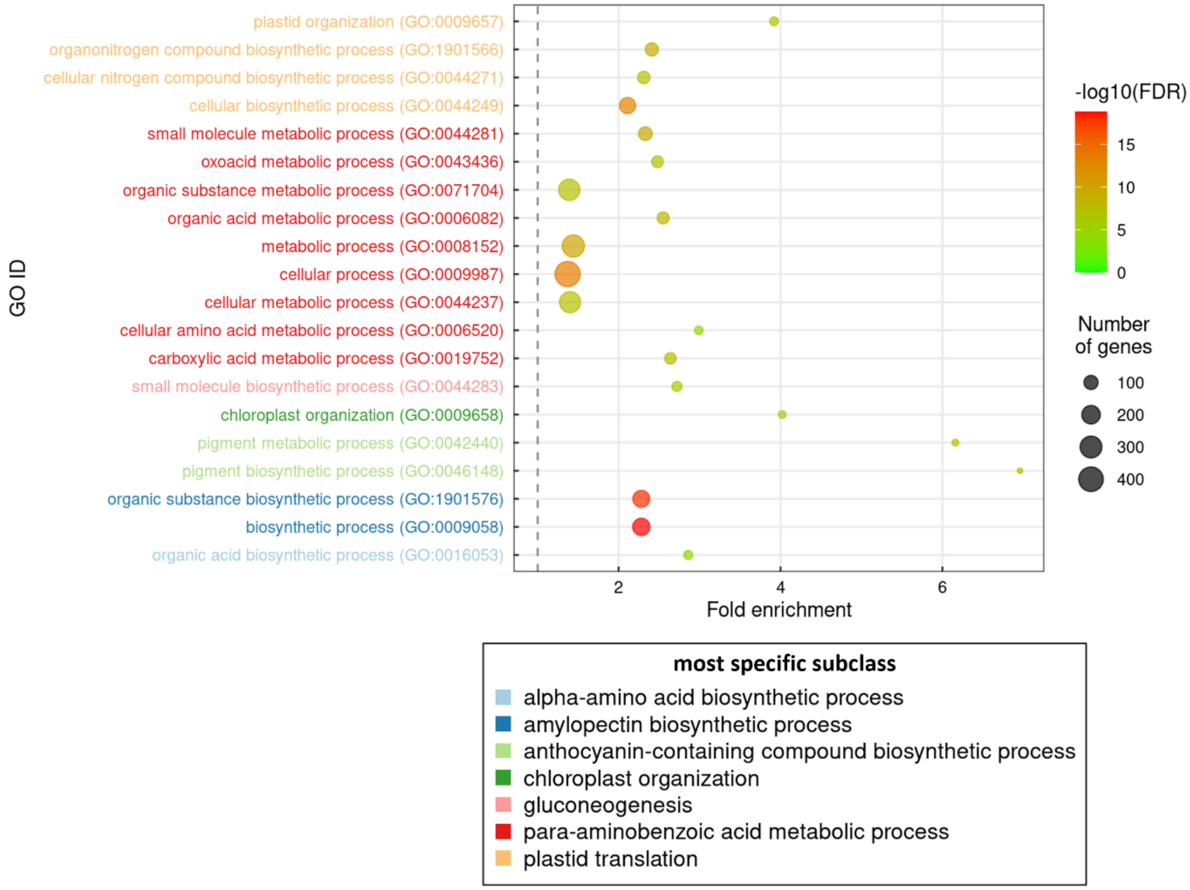

**Fig. S3** 6-day-old *lss* and *ess* are not impaired in the oxidative stress response. RT-qPCR analysis measuring the transcript abundance of the $^1O_2$ responsive genes *BAP1*, *ERF5*, *LOX3*, *AAA* and the $H_2O_2$ responsive genes *APX1*, *FER1*, *pKsi* at 6 days of growth. Bars indicate mean and lines indicate the standard error (n = 3 independent biological replicates). Statistical tests (pairwise t-test) shown against respective WT controls, * *P<0.05*.

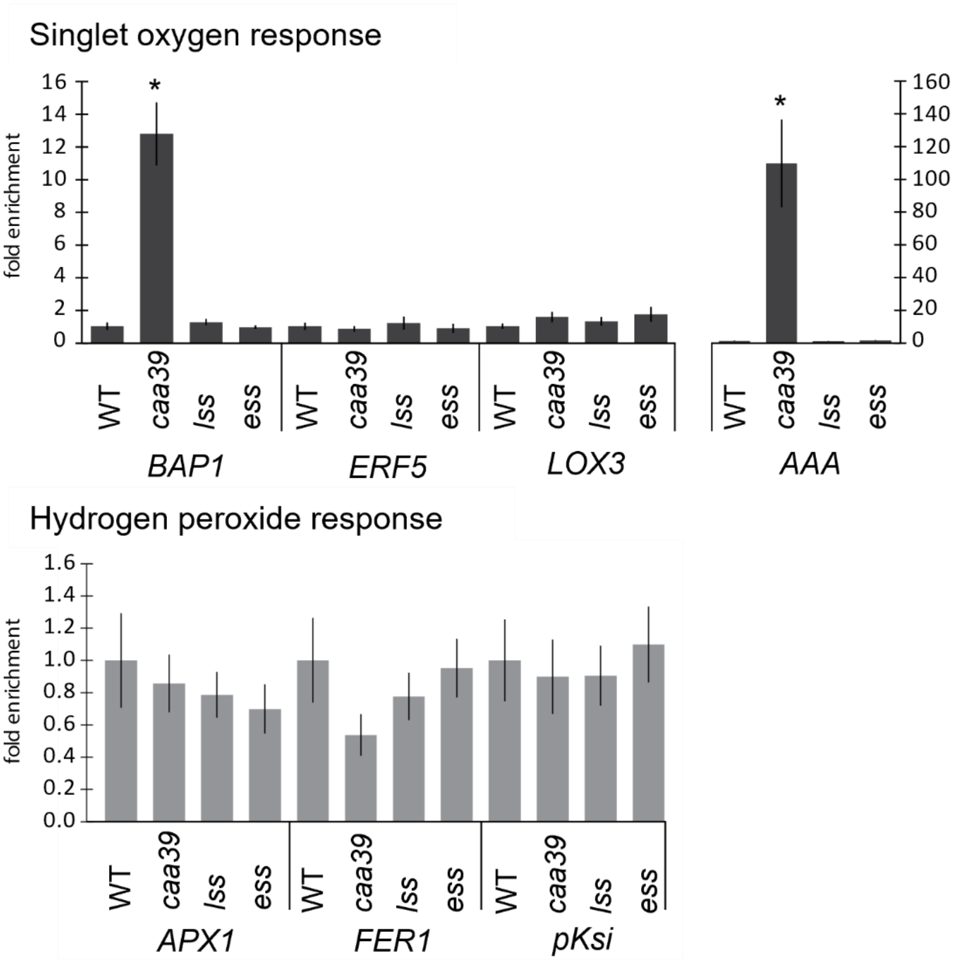


**Fig. S4** Suppressing AtSMC3 and AtTopoVI A leads to additive morphological phenotypes. Pictures of long day grown WT, *caa39, lss*, *caa39 lss*, *ess* and *caa39 ess* at 6 days, 2 weeks and 4

weeks. The pictures of WT, *caa39*, *lss* and *ess* are the same as in figure 2 (a). Scale bar, 1cm.

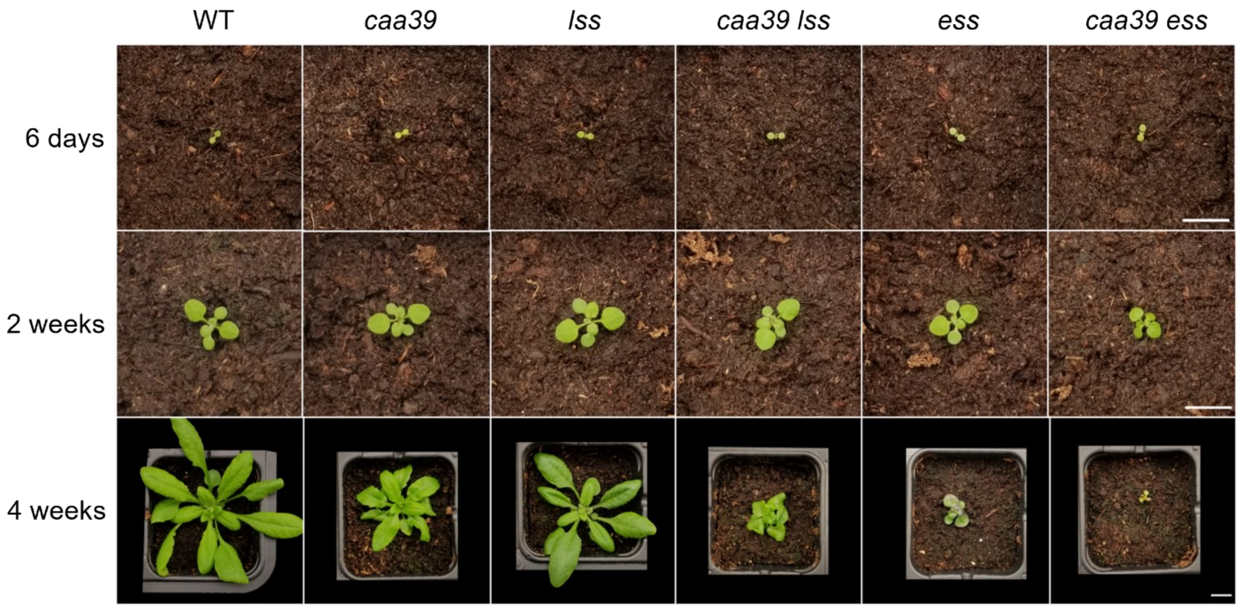


**Fig. S5** 4-week-old *lss* displays a slight tolerance to photooxidative stress. (a) Pictures of representative 4-week-old WT, *caa39*, *lss* and *caa39 lss* plantlets at 0 h, steady state; 32 h, right after the high-light exposure. Scale bar, 1cm. (b) Maximum PSII efficiency (Fv/Fm) of WT, *caa39*, *lss* and *caa39 lss* at 0 h and 32 h. Black lines indicate the standard error (n = 12 plants from 3 independent biological replicates). Compact letter display represents statistic groups and was generated using a pairwise t-test corrected with the Benjamini/Hochberg FDR method. (c) Non Photochemical Quenching (NPQ) measurements. Black lines indicate the standard error (n = 12 plants from 3 independent biological replicates). Compact letter display represents statistic groups and was generated using a pairwise t-test corrected with the

Benjamini/Hochberg FDR method.

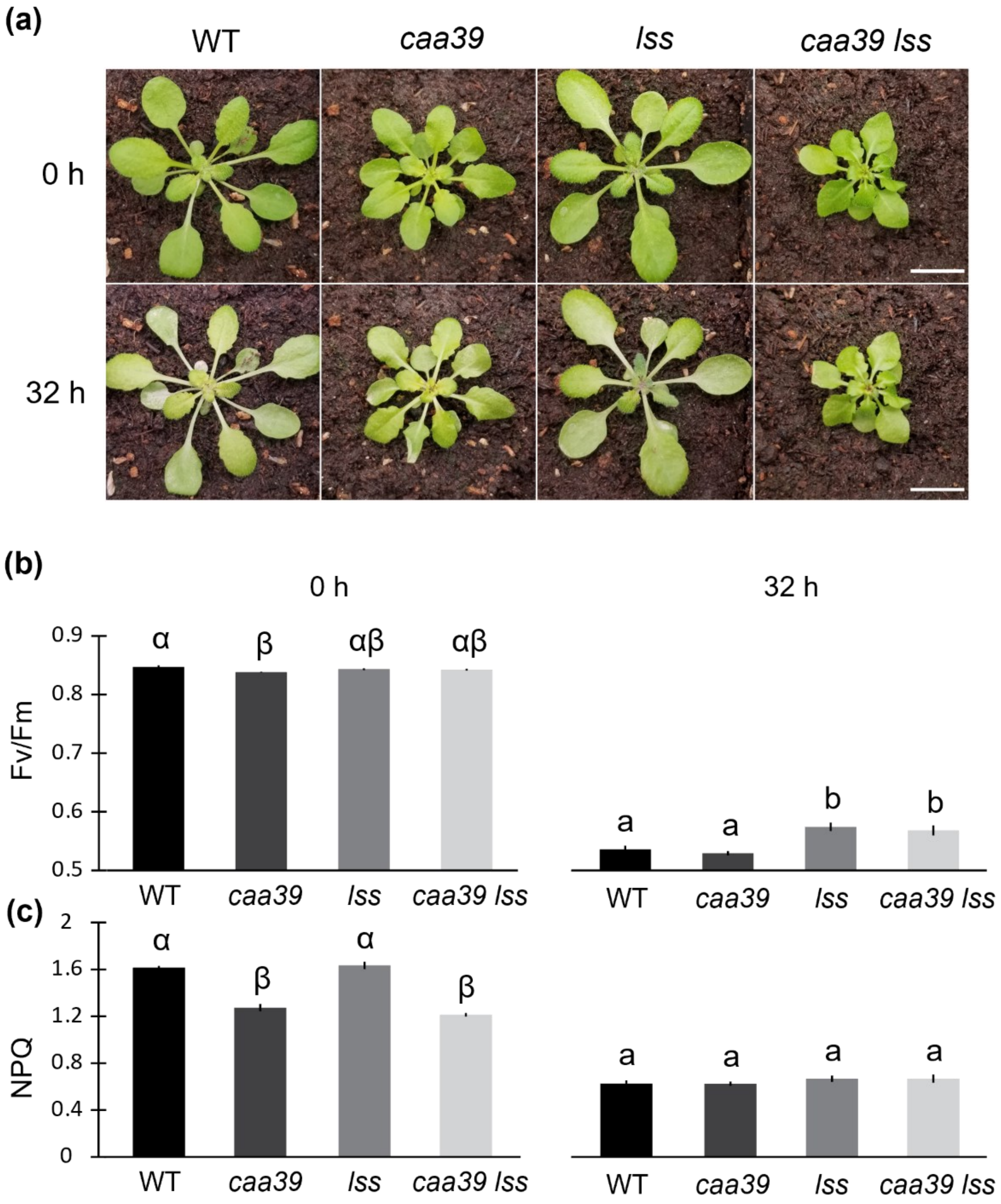
