## Supplemental Method 1 for "Structural Maintenance of Chromosome 3 interacts with the Topoisomerase VI complex and contributes to the oxidative stress response in *Arabidopsis thaliana*"

~~~~
---
title: 'Clustering of ARA2 data - 2 weeks plants'
author: "Cécile Lecampion"
date: "`r format(Sys.time(), '%m/%Y')`"
output:
  html_document:
    theme: cerulean
    highlight: tango
    df_print: paged
    toc: true
    toc_float: true
    number_sections: true
  pdf_document:
    toc: yes
    toc_float: true
    number_sections: yes
---
***

```{r setup, include=FALSE}
knitr::opts_chunk$set(echo = TRUE)
```
This script is derived from https://2-bitbio.com/2017/10/clustering-rnaseq-
data-using-k-means.html

### Data

This report analyses the data for the 2 weeks plants

<u>3 lines, 2 weeks</u>

WT

caa39

ess

```{r message=FALSE, echo=FALSE, warning= FALSE}
isRunOnMac <- Sys.info()[['sysname']] == 'Darwin'
WORKING_DIRECTORY <- ""
if (isRunOnMac) {
  WORKING_DIRECTORY <- "/Volumes/Disk_4To/Donnees_ARA2/count"
  setwd(WORKING_DIRECTORY)
} else {
  WORKING_DIRECTORY <- "~/partage/F/Donnees_ARA2/count"
  setwd(WORKING_DIRECTORY)
}

wt1_2w <- read.table("wt1_2w_sorted.count", header = FALSE, col.names =
c("Id", "wt1_2w"))
wt2_2w <- read.table("wt2_2w_sorted.count", header = FALSE, col.names =
c("Id", "wt2_2w"))
~~~~

````
caa1_2w <- read.table("caa39_1_2w_sorted.count", header = FALSE, col.names =
c("Id", "caa1_2w"))
caa2_2w <- read.table("caa39_2_2w_sorted.count", header = FALSE, col.names =
c("Id", "caa2_2w"))

ess1_2w <- read.table("ess_1_2w_sorted.count", header = FALSE, col.names =
c("Id", "ess1_2w"))
ess2_2w <- read.table("ess_2_2w_sorted.count", header = FALSE, col.names =
c("Id", "ess2_2w"))

filtre <- read.table("list_DE-gene_2w.txt", header = FALSE)
```
Raw reads count for all lines are gathered in a single dataframe.

```{r echo=FALSE, message= FALSE}
require(plyr)
count2w <- Reduce(function(x,y) merge(x = x, y = y, by = "Id"),
                list(wt1_2w, wt2_2w, caa1_2w, caa2_2w, ess1_2w, ess2_2w))
Hcount2w <- head(count2w)
knitr::kable(Hcount2w,
             caption = "First lines of the data frame ")

```
Counts are normalised in edgeR to take library size into account

```{r echo=FALSE, message= FALSE, warning = FALSE}
require(edgeR)
### Normalisation
y2w <- as.matrix(sapply(count2w, as.numeric))
y2w <- y2w[, colnames(y2w) != "Id"]
y2w <- DGEList(counts = y2w, group=c("wt1_2w", "wt2_2w", "caa1_2w", "caa2_2w",
"ess1_2w", "ess2_2w"))
y2w <- calcNormFactors(y2w)
z2w <- cpm(y2w, normalized.lib.size=TRUE)
row.names(z2w) <- count2w$Id

```


# Genes selection
Data are filtered to keep only genes that are either up or down regulated in
at least one pairwise comparaison : caa39 vs wt, ess vs wt et ess vs caa39
```{r echo=FALSE, message= FALSE, warning = FALSE}
library(dplyr)
z2w <- as.data.frame(z2w)
z2w <- z2w %>% filter(row.names(z2w) %in% filtre$V1)
```


```{r echo=FALSE, message= FALSE, warning = FALSE}
### Building a tree
z2w <- as.matrix(z2w)
scaledata2w <- t(scale(t(z2w)))
hc2w <- hclust(as.dist(1-cor(scaledata2w, method="spearman")),
method="complete")
````

````
TreeC2w = as.dendrogram(hc2w, method="average")
plot(TreeC2w,
main = "Sample Clustering",
ylab = "Height")

```
````

### How many clusters should be create ?

There are a few methods for evaluating the optimum number of clusters.

#### SSE : sum of squared error
SSE is defined as the sum of the squared distance between each member of a cluster and its cluster centroid. We repeatedly test and increasing number of clusters and evaluate the SSE. As we increase the number of clusters the distance between any point and it’s centroid will be smaller since the cluster itself is smaller. At a certain number of clusters number however, the SSE will not significantly decrease with each new addition of a cluster. This is the elbow and suggests a suitable number of clusters:

````
```{r echo=FALSE, message= FALSE, warning = FALSE}
wss2w <- (nrow(scaledata2w)-1)*sum(apply(scaledata2w,2,var))
for (i in 2:20) wss2w[i] <- sum(kmeans(scaledata2w,
centers=i)$withinss)
plot(1:20, wss2w, type="b", xlab="Number of Clusters",
ylab="Within groups sum of squares")

```
````

#### Average silhouette width
The next method is by estimating the optimum number using the average silhouette width. The silhouette value describes how similar a gene is to its own cluster (cohesion) compared to other clusters (separation). A high value indicates that the gene is well placed. So if the average of all of these silhouettes is high then the number of clusters is good.

````
```{r echo=FALSE, message= FALSE, warning = FALSE}
library(cluster)
sil2w <- rep(0, 20)
for(i in 2:20){
k1to202w <- kmeans(scaledata2w, centers = i, nstart = 25, iter.max = 20)
ss2w <- silhouette(k1to202w$cluster, dist(scaledata2w))
sil2w[i] <- mean(ss2w[, 3])
}
plot(1:20, sil2w, type = "b", pch = 19, xlab = "Number of clusters k",
ylab="Average silhouette width")
abline(v = which.max(sil2w), lty = 2)
#cat("Average silhouette width optimal number of clusters:", which.max(sil),
"\n")
```
````
<br>
`r {"Average silhouette width optimal number of clusters : "}`
`r which.max(sil2w)`
````

### Calinsky criterion
The Calinski-Harabasz index is based on the intra and inter cluster sum of squares. So we are looking to maximize the index to find well separated clusters.

```{r echo=FALSE, message= FALSE, warning = FALSE}
library(vegan)
fit2w <- cascadeKM(scaledata2w, 1, 20, iter = 100)
plot(fit2w, sortg = TRUE, grpmts.plot = TRUE)
calinski.best2w <- as.numeric(which.max(fit2w$results[2,]))
```
<br>
`r {"Calinski criterion optimal number of clusters : "}`
`r calinski.best2w`

## Gap statistic
Next up is the gap statistic. The gap statistic comapres the log within-cluster sum of squares (discussed above) with it’s expectation under the null reference distribution. Then it chooses the cluster where the gap between the log(wss) and the maximim of the null ref is the largest:

```{r echo=FALSE, message= FALSE, warning = FALSE}
library(cluster)
set.seed(13)
gap2w <- clusGap(scaledata2w, kmeans, 20, B = 100, verbose = interactive())
plot(gap2w, main = "Gap statistic")
abline(v=which.max(gap2w$Tab[,3]), lty = 2)
```

```{r echo=FALSE, message= FALSE, warning = FALSE}
GENE_CLUSTER_SCORE <- 0.85
CLUSTER_COUNT <- 7
```

Select `r which.max(sil2w)` clusters lead to oversimplification of the data, `r which.max(gap2w$Tab[,3])` clusters is probably to large. We have to select a value between those two. let's select `r CLUSTER_COUNT` clusters.

# Clustering the data

For more readable plot, we use the mean of the 2 replicates for each line.
```{r echo=FALSE, message= FALSE, warning = FALSE}
### Mean of the column grouped by line
z2w_mean <- as.matrix(rowMeans(z2w[, 1:2]))
z2w_mean <- cbind(z2w_mean, (rowMeans(z2w[, 3:4])))
z2w_mean <- cbind(z2w_mean, (rowMeans(z2w[, 5:6])))
colnames(z2w_mean) <- c("wt", "caa39", "ess")
scaledata_mean <- t(scale(t(z2w_mean)))
```

## Plotting the centroids to show behavior
```{r echo=FALSE, message= FALSE, warning = FALSE}
#clustering
set.seed(20)
```

````
kClust <- kmeans(scaledata_mean, centers=CLUSTER_COUNT, nstart = 1000,
iter.max = 20)
kClusters <- kClust$cluster

#find centroid
clust.centroid = function(i, dat, clusters) {
ind = (clusters == i)
colMeans(dat[ind,])
}

kClustcentroids <- sapply(levels(factor(kClusters)), clust.centroid,
scaledata_mean, kClusters)
library(ggplot2)
library(reshape)
#get in long form for plotting
Kmolten <- melt(kClustcentroids)
colnames(Kmolten) <- c('sample','cluster','value')
p <- ggplot(Kmolten, aes(x=sample,y=value, group=cluster,
colour=as.factor(cluster))) +
geom_point() +
geom_line() +
xlab("Line") +
ylab("Expression") +
labs(title= "Cluster Expression by Line",color = "Cluster")
p
```

## How similar the clusters are ?

Computing the correlation between centroids, show the similarity of the
clusters between each otehr.If the correlation between 2 clusters is to high,
let say > 0.85, then the number of clusters should be lowered.

`r {knitr::kable(cor(kClustcentroids), row.names = TRUE)}`

To check clusters quality (are the gene correctly assigned ?) we can plot the
behavior of each gene in a cluster.

# Clusters analysis

```{r echo=FALSE, message= FALSE, warning = FALSE}

#------------------------------------------------------------------------------------------
### Plot the behavior of the genes belonging to cluster number "clusterNb"
### compare to the cluster centroïd
### Return a list:
### - table: Table named Kmolten with ID and a correlation score
### - plot: the plot
#------------------------------------------------------------------------------------------
f_etudeCluster <- function(clusterNb, Kmolten) {

  core <- Kmolten[Kmolten$cluster == clusterNb,]
  K <- (scaledata_mean[kClusters == clusterNb,])
  corscore <- function(x){cor(x,core$value)}
````

```
  score <- apply(K, 1, corscore)
  KmoltenC <- melt(K)
  colnames(KmoltenC) <- c('gene','sample','value')
  #add the score
  KmoltenC <- merge(KmoltenC,score, by.x='gene',by.y='row.names', all.x=T)
  colnames(KmoltenC) <- c('gene','sample','value','score')

  # To plot only a list of gene and if the "filter list" doesn't contain
transcript number
  # library(dplyr)
  # library(stringr)
  # KmoltenC %>% filter(str_detect(gene, str_c(oxrep$V1, collapse="|")))

  #order the dataframe by score
  #to do this first create an ordering factor
  KmoltenC$order_factor <- 1:length(KmoltenC$gene)
  #order the dataframe by score
  KmoltenC <- KmoltenC[order(KmoltenC$score),]
  #set the order by setting the factors
  KmoltenC$order_factor <- factor(KmoltenC$order_factor , levels =
KmoltenC$order_factor)
  # Everything on the same plot
  p <- ggplot(KmoltenC, aes(x=sample,y=value)) +
    geom_line(aes(colour=score, group=gene)) +
    scale_colour_gradientn(colours=c('blue1','red2')) +
    #this adds the core
    geom_line(data=core, aes(sample,value, group=cluster),
color="black",inherit.aes=FALSE) +
    xlab("Line") +
    ylab("Expression") +
    labs(title= sprintf("Cluster %d Expression", clusterNb),color = "Score")
  # Retourne le tableau et le graphe
  return(list(table=KmoltenC, plot=p))
}

### Getting the gene ID for further anlysis (only if score > GENE_CLUSTER_SCORE)
### Dataframe to collect the number of gene in each cluster before and after
filtering on score
df_cluster_length <- data.frame(matrix(ncol = 3, nrow = 0))
colnames(df_cluster_length) <- c("raw", "filtered", "retained")

library(tidyverse)

for (clusterNb in c(1:CLUSTER_COUNT)) {
  etude <- f_etudeCluster(clusterNb, Kmolten)
  print(etude$plot)
  KmoltenC <- etude$table
  cluster <- KmoltenC$gene[KmoltenC$score> GENE_CLUSTER_SCORE]
  cluster <- as.data.frame(cluster[!duplicated(cluster)])
  colnames(cluster) <- sprintf("cluster%d", clusterNb)
  df_cluster_length <- df_cluster_length %>% add_row(raw =
as.numeric(length(unique(KmoltenC$gene))), filtered =
as.numeric(nrow(cluster)))
```

  write.table(cluster, file = sprintf("%s/cluster%d.txt", WORKING_DIRECTORY,
clusterNb), quote = FALSE, row.names = FALSE)
}

df_cluster_length$retained <- round(df_cluster_length$filtered/
df_cluster_length$raw*100, digits = 2)
```

Gene Id for each cluster are exported to proceed to Gene Ontology analysis.
Only the genes with a score > `r GENE_CLUSTER_SCORE` are kept.

&lt;br&gt;

`r knitr::kable(df_cluster_length,caption = "% of gene retained after
filtering on score", row.names = TRUE)`

### Gene ontology
The aim is to find gene ontology enrichment among the differentially expressed
genes.
Preparation of the data is performed by the `prepare_gene_ontology.pl` script
(Terese et Lecampion : https://github.com/cecile-lecampion/gene-ontology-
analysis-and-graph). It uses PANTHER and REVIGO.
Colors link each GO biological process to the main process they belong.

```{zsh engine.opts="-l", include=FALSE}
#Préparation des données
if [ "$(uname -s)" = 'Darwin' ]; then
  cd /Volumes/Disk_4To/Donnees_ARA2
else
  cd ~/partage/F/Donnees_ARA2
fi

# Determine the number of cluster based on the number of files cluster*.txt in
the directory ./count
CLUSTER_COUNT=$(ls -1 count/cluster*.txt | wc -l)

# Perform GO for each cluster
for CLUSTER_NB in $(seq 1 $CLUSTER_COUNT); do
  ~/mytools/prepare_gene_ontology.pl --method biological_process --correction
fdr \
                                     ./count/cluster${CLUSTER_NB}.txt \
                                     ./cluster${CLUSTER_NB}
_curated_gene_ontology.tsv \
                                     ./cluster${CLUSTER_NB}
_go_ids_hierarchy.tsv
done
```

```{r echo=FALSE, message=FALSE, warning=FALSE, fig.width = 9}

isRunOnMac <- Sys.info()[['sysname']] == 'Darwin'
if (isRunOnMac) setwd("/Volumes/Disk_4To/Donnees_ARA2") else
  setwd("~/partage/F/Donnees_ARA2")

```
#-----------------------------------------------------------------------------------------
### Perform Gene Ontology for the cluster clusterNb
#-----------------------------------------------------------------------------------------
f_determineGeneOntology <- function(clusterNb) {
  tsvFile <- sprintf("cluster%d_curated_gene_ontology.tsv", clusterNb)
  GO <- read.table(tsvFile, header=T, stringsAsFactors = T, sep = "\t")
  GO$GO_id <-gsub("biological_process_involved_in_","",as.character(GO$GO_id))

  hierarchytsvfile <- sprintf("cluster%d_go_ids_hierarchy.tsv", clusterNb)
  hierarchy <- read.table(hierarchytsvfile, header=F, stringsAsFactors = T,
sep = "\t", quote = "")
  colnames(hierarchy) <- c("parent", "child")
  hierarchy$child <-gsub("biological process involved in
","",as.character(hierarchy$child))

  # If you only want to use the n first line of the data frame for the plot,
execute this command
  # If you want to keep all lines just skip this
  #Data in the input file are sorted in ascending FDR.

  GO <- GO[1:20,]  #Replace the data frame by a new data frame that only
contains the 20 first lines of GO
                    # and all columns
                    # You can select any number of lines to be used by
replacing 20 by the desired value

  # List objects and their structure contained in the dataframe 'GO'
  ls.str(GO)

  # Transform the column 'Gene_number' into a numeric variable
  GO$Gene_number <- as.numeric(GO$Gene_number)

  # Replace all the "_" by a space in the column containing the GO terms
  GO$GO_id <- chartr("_", " ", GO$GO_id)

  # Transform FDR values by -log10('FDR values')
  GO$'|log10(FDR)|' <- -(log10(GO$FDR))

  # Add parent GO-ID to GO
  parent <- hierarchy[hierarchy$child %in% GO$GO_id, ]
  # all = TRUE to keep all the lines
  GO <- merge(GO, parent, by.x=c("GO_id"), by.y=c("child"), all = TRUE)
  # Convert column GO$parent to character (it is factor) to replace tha NA by
the corresponding value of column GO_id
  GO$parent <- as.character(GO$parent)
  # Suppress the text between ()
  GO[is.na(GO)] <- gsub("\\s*\\([^\\)]+\
\)","",as.character(GO$GO_id[is.na(GO$parent)]))
  # Make GO$parent factor again
  GO$parent <- factor(GO$parent)
  GO <- GO[order(GO$parent),]

  # Prepare color for parent GO-Id assignment
```

```
  numColors <- length(levels(GO$parent))
  getColors <- scales::brewer_pal('qual', palette = "Paired")
  myPalette <- getColors(numColors)
  names(myPalette) <- levels(GO$parent)

  # Draw the plot with ggplot2
  #--------------------------------------
  p <- ggplot(GO, aes(x = GO_id, y = Fold_enrichment)) +
      geom_hline(yintercept = 1, linetype="dashed",
                 color = "azure4", size=.5)+
      geom_point(data=GO, aes(x=GO_id, y=Fold_enrichment,
                               size = Gene_number, colour = `|log10(FDR)|
`), alpha=.7)+
      # scale_y_continuous(limits = c(0,15))+
      scale_x_discrete(limits= GO$GO_id)+
      scale_color_gradient(low="green", high="red", limits=c(0, NA))+
      coord_flip()+
      theme_bw()+
      theme(axis.ticks.length=unit(-0.1, "cm"),
            axis.text.x = element_text(margin=margin(5,5,0,5,"pt"), color =
"black"),
            axis.text.y = element_text(margin=margin(5,5,5,5,"pt"),
colour=myPalette[GO$parent]),
            #axis.text = element_text(color = "black"),
            panel.grid.minor = element_blank(),
            legend.title.align=0.5)+
      xlab("GO ID")+
      ylab("Fold enrichment")+
      ggtitle(sprintf("cluster%d", clusterNb))+
      # Replace by your variable names; \n allow a new line for text
      labs(color="-log10(FDR)", size="Number\nof genes")+
      guides(size = guide_legend(order=2),
             colour = guide_colourbar(order=1))

  print(p)

  plot(NULL, xlim=c(0,length(myPalette)), ylim=c(0,1), xlab="", ylab="",
xaxt="n", yaxt="n", frame.plot = FALSE)
  legend("center", title = sprintf("cluster%d main GO-Id", clusterNb), legend
= names(myPalette), col = as.data.frame(myPalette)$myPalette, pch = 15, cex=1,
pt.cex = 1.5)

}

for (clusterNb in c(1:CLUSTER_COUNT)) {
  f_determineGeneOntology(clusterNb)
}
```

# Bibliography

T. Bonnot, MB. Gillard and DH. Nagel. "A Simple Protocol for Informative Visualization of Enriched Gene Ontology Terms". Bio-101: e3429. DOI:10.21769/BioProtoc.3429

Chen, Yunshun, Aaron A T Lun, and Gordon K Smyth. 2016. “From Reads to Genes to Pathways: Differential Expression Analysis of RNA-Seq Experiments Using Rsubread and the edgeR Quasi-Likelihood Pipeline.” F1000Research 5: 1438. https://doi.org/10.12688/f1000research.8987.2.

Huaiyu Mi, Dustin Ebert, Anushya Muruganujan, Caitlin Mills, Laurent-Philippe Albou, Tremayne Mushayamaha and Paul D Thomas. 2020. "PANTHER version 16: a revised family classification, tree-based classification tool, enhancer regions and extensive API". Nucl. Acids Res. (2020) doi: 10.1093/nar/gkaa1106s.

Maechler, Martin, Peter Rousseeuw, Anja Struyf, Mia Hubert, and Kurt Hornik. 2021. Cluster: Cluster Analysis Basics and Extensions. https://CRAN.R-project.org/package=cluster.

Oksanen, Jari, F. Guillaume Blanchet, Michael Friendly, Roeland Kindt, Pierre Legendre, Dan McGlinn, Peter R. Minchin, et al. 2020. Vegan: Community Ecology Package. https://CRAN.R-project.org/package=vegan.

R Core Team. 2021. R: A Language and Environment for Statistical Computing. Vienna, Austria: R Foundation for Statistical Computing. https://www.R-project.org/.

Supek F, Bošnjak M, Škunca N, Šmuc T. "REVIGO summarizes and visualizes long lists of Gene Ontology terms" .PLoS ONE 2011. doi:10.1371/journal.pone.0021800

Wickham, Hadley. 2007. “Reshaping Data with the Reshape Package.” Journal of Statistical Software 21 (12). http://www.jstatsoft.org/v21/i12/paper.

Wickham, Hadley. 2011. “The Split-Apply-Combine Strategy for Data Analysis.” Journal of Statistical Software 40 (1): 1–29. http://www.jstatsoft.org/v40/i01/.

Wickham, Hadley. 2016. Ggplot2: Elegant Graphics for Data Analysis. Springer-Verlag New York. https://ggplot2.tidyverse.org.

Wickham, Hadley, Mara Averick, Jennifer Bryan, Winston Chang, Lucy D’Agostino McGowan, Romain François, Garrett Grolemund, et al. 2019. “Welcome to the tidyverse.” Journal of Open Source Software 4 (43): 1686. https://doi.org/10.21105/joss.01686.

Wickham, Hadley, Romain François, Lionel Henry, and Kirill Müller. 2022. Dplyr: A Grammar of Data Manipulation. https://CRAN.R-project.org/package=dplyr.

Wickham, Hadley, and Dana Seidel. 2020. Scales: Scale Functions for Visualization. https://CRAN.R-project.org/package=scales.

Xie, Yihui. 2014. “Knitr: A Comprehensive Tool for Reproducible Research in R.” In Implementing Reproducible Computational Research, edited by Victoria Stodden, Friedrich Leisch, and Roger D. Peng. Chapman; Hall/CRC. http://www.crcpress.com/product/isbn/9781466561595.

Xie, Yihui, Christophe Dervieux, and Emily Riederer. 2020. R Markdown Cookbook. Boca Raton, Florida: Chapman; Hall/CRC. https://bookdown.org/yihui/rmarkdown-cookbook.

Script prepare_gene_onthology.pl. Terese M. et Lecampion C. https://github.com/cecile-lecampion/gene-ontology-analysis-and-graph

# R session information and parameters

```{r echo=FALSE, message=FALSE, warning=FALSE}

devtools::session_info()
```
